# APOE traffics to astrocyte lipid droplets and modulates triglyceride saturation and droplet size

**DOI:** 10.1101/2023.04.28.538740

**Authors:** Ian A. Windham, Joey V. Ragusa, E. Diane Wallace, Colby H. Wagner, Kristen K. White, Sarah Cohen

**Affiliations:** Department of Cell Biology and Physiology, University of North Carolina at Chapel Hill; Mass Spectrometry Core Laboratory, Department of Chemistry, University of North Carolina at Chapel Hill; Microscopy Services Laboratory, Department of Pathology and Laboratory Medicine, University of North Carolina at Chapel Hill

**Author notes:** Correspondences: Sarah Cohen.

## Abstract

The *E4* variant of *APOE* strongly predisposes individuals to late-onset Alzheimer’s disease. We demonstrate that in response to neutral lipid synthesis, apolipoprotein E (APOE) in astrocytes can avoid translocation into the ER lumen and traffic to lipid droplets (LDs) via membrane bridges at ER-LD contacts. *APOE* knockdown promotes fewer, larger LDs containing more unsaturated triglyceride. This LD size distribution phenotype was rescued by chimeric APOE that targets only LDs. APOE4*-*expressing astrocytes also form a small number of large LDs enriched in unsaturated triglyceride. Additionally, the larger LDs in *APOE4* cells exhibit impaired turnover and increased sensitivity to lipid peroxidation. Our data indicate that APOE plays a previously unrecognized role as an LD surface protein that regulates LD size and composition.

*APOE4* is a toxic gain of function variant that causes aberrant LD composition and morphology. We propose that *APOE4* astrocytes with large, unsaturated LDs are sensitized to lipid peroxidation or lipotoxicity, which could contribute to Alzheimer’s disease risk.

**Summary:** Windham *et al*. discover that APOE in astrocytes can traffic to lipid droplets (LDs), where it modulates LD composition and size. Astrocytes expressing the Alzheimer’s risk variant APOE4 form large LDs with impaired turnover and increased peroxidation sensitivity.

## Introduction

Lipids comprise 60% of the brain’s dry mass (O’Brien and Sampson, 1965). Consequently, sophisticated molecular mechanisms evolved to manage the distribution and utilization of the diverse lipid species present in the brain. Astrocytes coordinate many aspects of brain lipid homeostasis; they mediate lipid uptake from the blood, synthesize lipids for neurons such as cholesterol and polyunsaturated fatty acids, and take up peroxidated lipids from neurons for detoxification (Pfrieger and Ungerer, 2011; Lee et al., 2021; Ralhan et al., 2023). Cytoplasmic lipid droplets (LDs) are pivotal components of astrocyte lipid homeostasis. LDs store lipids including triglycerides (TG) and cholesterol esters (CE) in a neutral lipid core surrounded by an amphipathic phospholipid monolayer. Proteins that coat the surface of LDs regulate their biogenesis and turnover, as well as a medley of other cellular processes including cell signaling, protein homeostasis, and inflammation. LDs store energy in the form of fatty acids that can be beta-oxidized, and also buffer against lipotoxic stress by preventing the accumulation of harmful lipid intermediates (Olzmann and Carvalho, 2019). Therefore, processes that regulate LD biogenesis and turnover are critical to protecting cells from lipid-related insults. Notably, Alois Alzheimer first observed the accumulation of “adipose inclusions” in the glia of patient brain tissue in his foundational study which also describes extracellular amyloid beta plaques and tau neurofibrillary tangles (Alzheimer, 1907; translated in Alzheimer et al., 1995). More recent work has demonstrated that both astrocytes and microglia accumulate LDs during aging and in pathologies such as ischemia, neuroinflammation, and neurodegenerative diseases including Alzheimer’s Disease (Farmer et al., 2020; Ralhan et al., 2023). However, it is unclear whether LD accumulation is a cause or consequence of pathology.

A key protein mediator of brain lipid homeostasis is apolipoprotein E (APOE), a 34 kDa secreted protein expressed primarily by astrocytes and microglia as well as neurons under stress (Xu et al., 2006). APOE is a component of high-density lipoprotein particles that transport lipids between cells in the brain. Nascent APOE lipoproteins are assembled in the lumen of endoplasmic reticulum (ER) prior to secretion, although the mechanism for this stage of biogenesis is unknown. After secretion, APOE-coated lipoproteins are lipidated via reverse cholesterol transport from the plasma membrane mediated by ABC transporters, primarily ABCA1. APOE on lipoproteins binds to cell-surface lipoprotein receptors, including LDLR, LRP, and APOER2, triggering lipoprotein uptake via receptor-mediated endocytosis (Hauser et al., 2011; Mahley, 2016).

Astrocyte-derived lipoproteins supply key lipids, predominately cholesterol and phospholipids, to neurons for building their high surface area membranes and cholesterol-rich lipid nanodomains at synapses (van Deijk et al. 2017). Other studies demonstrated that lipid peroxides formed in neurons can be transported by APOE-lipoproteins to astrocytes, which may play a role in detoxifying these deleterious lipid species (Liu et al., 2017; Ioannou et al., 2019). *APOE* is notably the strongest genetic risk factor for late-onset Alzheimer’s disease. The *APOE4* variant is a cysteine to arginine substitution at residue 112. Individuals possessing the *APOE4* variant are significantly more likely to develop late-onset Alzheimer’s disease than those homozygous for the *APOE3* variant in a dose-dependent manner (Corder et al., 1993). There is no consensus mechanism for how the *E4* variant predisposes individuals to Alzheimer’s disease.

Although early work focused on the effects of *APOE4* on the clearance of amyloid-β, a number of recent studies draw a compelling connection between *APOE4* and glia-specific lipid dishomeostasis. Multiple transcriptomic studies of *APOE4*-expressing iPSC-derived astrocytes and microglia demonstrate alterations in the expression of lipid metabolic genes as well as defects in cholesterol trafficking and metabolism (Lin et al. 2018; de Leeuw et al. 2022; Tcw et al. 2022). Another study demonstrated that iPSC-derived *APOE4* astrocytes accumulate LDs rich in highly unsaturated TGs. Blocking fatty acid desaturation through inhibition of stearoyl-CoA desaturase 1, an enzyme elevated in AD patient brains, or supplementing cells with choline reduced LD accumulation in *APOE4* iPSCs (Astarita et al., 2011; Sienski et al. 2021)*. APOE4* iPSCs also exhibit defects in endocytosis (Narayan et al. 2020). Mouse astrocytes expressing human APOE4 have defects in LD metabolism as well as fatty acid uptake and oxidation (Farmer et al. 2019; Qi et al. 2021). However, it is unclear whether these phenotypes are downstream of APOE-mediated lipid secretion and uptake or another function of APOE within the cell.

Hints that APOE may play roles beyond lipid secretion came from studies of the LD surface proteome. A proximity ligation strategy identified APOE as a protein on the cytoplasmic face of LDs in Huh7 hepatocarcinoma cells (Bersuker et al. 2018). To be a positive hit, the protein must be accessible to APEX2-tagged PLIN2 on the surface of LDs, significantly reducing the possibility that this was a false positive caused by contamination of ER membranes in the LD preparation. This suggested that in addition to being secreted, APOE also has the capacity to target LDs in the cytoplasm. APOE was identified in subsequent studies of the LD proteome in mouse liver and THP-1 macrophages (Krahmer et al., 2018; Mejhert et al. 2020). Studies on the trafficking of APOE have also suggested that the cleaved C-terminus of APOE can exist in the cytoplasm and associate with mitochondria in neurons (Chang et al. 2005). Other members of the exchangeable apolipoprotein family have also been found to associate with LDs. APOE and APOCIII were identified on nuclear LDs in HepG2 cells, and APOAV has previously been reported on the surface of cytoplasmic LDs in adipocytes (Shu et al., 2010; Gao et al., 2012; Sołtysik et al. 2019). A putative LD-associated pool of APOE within glial cells could play an important role maintaining lipid homeostasis in the brain.

We sought to determine whether APOE is a *bona fide* LD protein in astrocytes. We observed that APOE localized to the cytoplasmic surface of LDs in astrocytes under conditions of neutral lipid synthesis. APOE trafficked to LDs by avoiding translocation into the endoplasmic reticulum (ER), targeting the cytoplasmic side of the ER before moving onto LDs at membrane bridges between the ER and LDs. By utilizing an oleic acid (OA) pulse-chase assay, we found that LD-associated APOE regulates the size of LDs. Knockdown of APOE caused a smaller number of large LDs enriched in unsaturated triglycerides to form during lipogenesis. Expression of an exclusively LD-targeted APOE chimeric construct rescued this phenotype, supporting a physiological role for APOE on the LD surface. Like APOE knockdown cells, APOE4-expressing cells had larger LDs than E3-expressing cells after OA pulse-chase, suggesting a lipid turnover defect. E4 LDs were dramatically enriched in unsaturated triglycerides and were more sensitive to lipid peroxidation than E3 LDs. We hypothesize these lipid homeostatic defects in E4 could sensitize astrocytes to stress and promote a reactive, proinflammatory state that contributes to Alzheimer’s pathogenesis.

## Results

### APOE localizes to cytoplasmic surface of LDs

Based on previous proteomics studies identifying APOE as a putative LD protein in liver and macrophages (Bersuker et al. 2018; Krahmer et al., 2018; Mejhert et al. 2020), we hypothesized that APOE localizes to LDs in astrocytes. Overexpressing a protein with lipid binding moieties could cause it to mislocalize to LDs. Therefore, it is imperative that LD-targeting of a protein is confirmed by probing its endogenous localization. To look at endogenous APOE in astrocytes, we employed immortalized *APOE3* targeted replacement astrocytes (TRAE3-H). These cells are astrocytes isolated from *APOE3* targeted replacement mice – a model in which the endogenous mouse coding exon is replaced by the human *APOE3* coding exon via homologous recombination – and subsequently immortalized via stable transfection of SV40 (Sullivan et al., 1997; Morikawa et al., 2005).

We stained fixed TRAE3-H cells for human APOE, LDs, and a marker of the Golgi. siRNA-mediated knockdown demonstrated the specificity of the APOE antibody (Fig. S1A-B). In complete media (CM), TRAE3-H cells contained few LDs, and APOE localized largely to Golgi apparatus and other endomembrane compartments, as expected. To model a LD-accumulating state, we loaded TRAE3h cells with 400 µM OA for 5 hours, which stimulates LD formation. Surprisingly, APOE relocalized to the surface of cytoplasmic LDs in a majority of cells after OA induction (Fig. 1A-C). We observed a concomitant reduction in colocalization of APOE with the Golgi marker GM130 (Fig. 1D). We also saw an increase in both intracellular APOE protein and, paradoxically, the amount of APOE protein secreted into the media upon OA treatment (Fig. S1C-E). We hypothesize that OA stimulates global APOE upregulation, and the subset of OA-treated cells which lack LD-associated APOE secrete more APOE than untreated cells.

**Figure 1:**
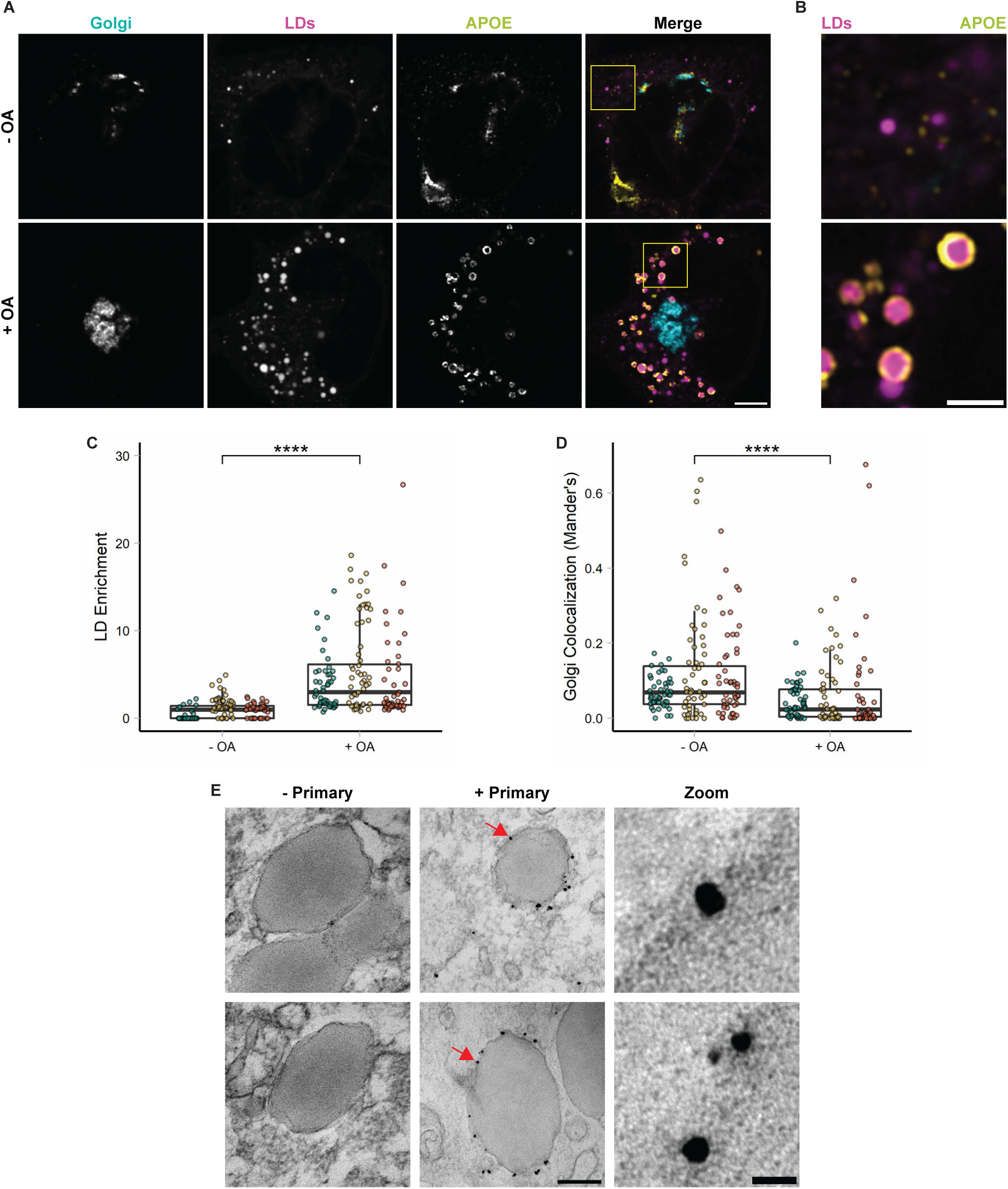
APOE localizes to cytoplasmic LDs in astrocytes during lipogenesis. (**A**) Representative confocal slices of TRAE3-H cells untreated (-OA) or treated with 400 µM oleate for 5 hours (+ OA). Cells were then fixed and stained for endogenous APOE with an anti-APOE antibody, the Golgi with an anti-GM130 antibody, and LDs with BODIPY 493/507. In the merged image, APOE is yellow, GM130 is cyan, and LDs are magenta. Scale bar, 10 µm. **(B)** Inset of images from (A) highlighting APOE enrichment on the surface of LDs after OA. Scale bar, 1 µm. **(C)** Quantification of APOE enrichment on the surface of LDs in TRAE3-H cells ± OA. LD enrichment was calculated by dividing the mean intensity of APOE fluorescence on LDs by the mean intensity of the non-LD associated APOE fluorescence in the cell. N = 150 cells per condition, with 50 cells from each independent experiment. Each data point represents one cell, and each color represents data collected from a separate, independent experiment. **(D)** Quantification of colocalization of APOE with the Golgi marker GM130 as measured by the Mander’s coefficient in TRAE3-H cells ± OA. The Mander’s coefficient was calculated by dividing the area of overlap between APOE and GM130 masks by the total area of the APOE mask. N = 50 cells per condition per experiment. Each data point represents one cell, and each color represents data collected from a separate, independent experiment. **(E)** Immunogold electron micrographs of endogenous APOE in TRAE3-H cells treated with 400 µM OA for 5 hours. The primary APOE antibody was included in images labelled “+ Primary”, and not included in the negative control images labelled “-Primary”. Silver-enhanced gold particles localize directly to the surface of LDs at the interface between the LD monolayer surface and the cytoplasm. Scale bars: 200 nm (left), 20 nm for zoom (right). P-values calculated using a clustered Wilcoxon rank sum test via the Rosner-Glynn-Lee method. **** p<0.0001

To see if we could observe LD-localized APOE in primary cells, we transfected primary rat astrocytes with human APOE tagged with mEmerald at the C-terminus. We observed APOE localized to LDs in a subset of cells grown in astrocyte media in the absence of OA treatment (S2A-B). Transfected APOE also localized to LDs in U-2 OS osteosarcoma cells (Fig. S2C).

The ability of APOE to localize to cytoplasmic LDs is particularly striking, as APOE is normally a secreted protein and possesses a canonical N-terminal signal peptide that targets it for translocation into the ER lumen (Zannis et al., 1984). Prior studies have shown that proteins with LD binding moieties that are targeted to the ER lumen become enriched within the ER at ER-LD contact sites. Their localization pattern resembles LD surface proteins, but the protein is within the lumen of the ER (Mishra et al., 2016). Therefore, although APOE appeared enriched on the surface of LDs, it was not clear whether this signal represented APOE protein bound to the cytoplasmic LD surface or within the ER lumen at an ER-LD contact site. To distinguish between these two possibilities, we utilized immunogold electron microscopy, staining for APOE in TRAE3h cells. We observed gold particles dotting the cytoplasmic face of both LDs and the ER (Fig. 1E). Gold was observed on the LD surface in the absence of any associated ER membrane, suggesting that APOE is a *bona fide* LD protein.

To further verify the topology of LD-associated APOE, we performed fluorescence protease protection assays (FPPs) (Fig. 2A) (Lorenz et al. 2006). Primary cortical rat astrocytes expressing a fluorescent marker of the ER lumen and APOE-mEm or another fluorescently-tagged LD protein were first treated with digitonin.

**Figure 2:**
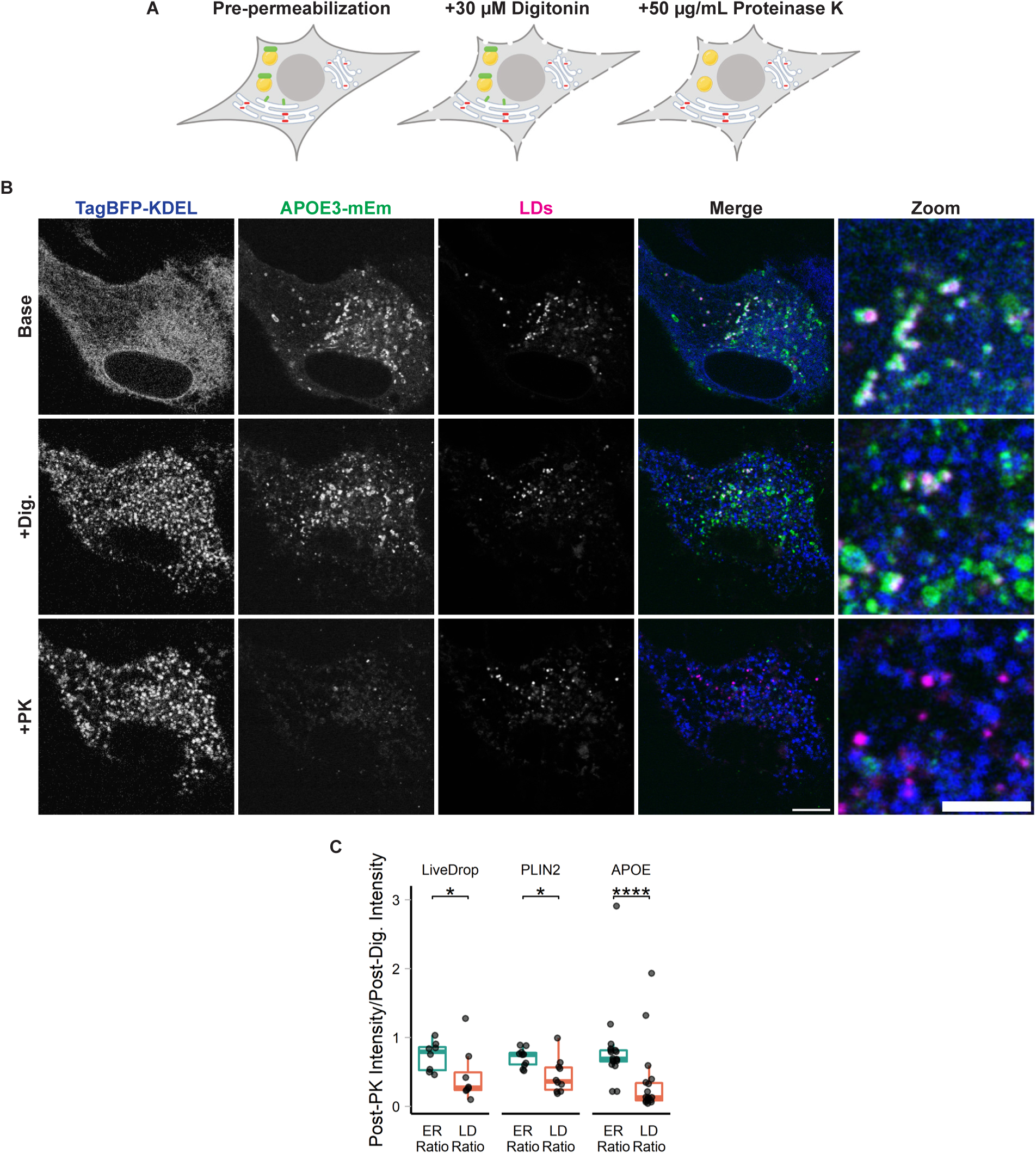
LD-associated APOE is exposed to the cytoplasm. (**A**) Cartoon schematic of the FPP assay to test the topology of fluorescently-tagged proteins in cells. Cells are treated with 30 µM digitonin for 1 minute, which selectively permeabilizes the plasma membrane but not the ER. After permeabilization, cells are treated with 50 µg/mL proteinase K, which enters the permeabilized plasma membrane and degrades all cytoplasmic-facing fluorophores (green). Because the ER membrane is not permeabilized, proteinase K does not enter into the ER lumen and ER lumen-facing fluorophores are retained (red). **(B)** Representative confocal slices of FPP performed on primary cortical rat astrocytes transiently transfected with APOE3-mEm, the ER marker TagBFP2-KDEL, and labelled for LDs with BODIPY 665/676. After digitonin permeabilization and proteinase K treatment, APOE signal on the surface of LDs is lost, but the luminal ER marker fluorescence is retained. A Gaussian filter with a radius of 1 pixel was applied to all images to improve visibility for print. Scale bars: 10 µm (left), 5 µm for zoom (right). **(C)** For APOE3 as well as two representative LD proteins, PLIN2 and LiveDrop, the fluorescence intensity ratio was calculated by dividing the mean fluorescence intensity after both permeabilization and proteinase K treatment by the mean fluorescence intensity after permeabilization. The ER ratio was calculated by measuring the whole cell intensity of the ER fluorescence at the two timepoints. The LD ratio was calculated by measuring the fluorescence on the surface of BODIPY-labelled LDs at the two timepoints. Ratios close to 1 indicate minimal loss of signal after proteinase K treatment, as observed with the ER marker TagBFP2-KDEL. Lower ratios indicate loss of fluorescence upon proteinase K treatment. N = 8-18 cells per condition, collected from three independent experiment. *p<0.05, **** p<0.0001. Dig., 30 µM digitonin. PK, +50 µg/mL Proteinase K. P-values were calculated via the Wilcoxon rank sum test and Bonferonni-corrected for multiple comparisons.

Digitonin selectively permeabilizes the plasma membrane while leaving the ER membrane intact. After digitonin permeabilization, proteinase K was added to degrade cytoplasmic-facing fluorophores while leaving ER-luminal proteins intact. Upon addition of proteinase K, the signal of the ER luminal marker was retained, while LD-associated proteins, including APOE, were degraded (Fig. 2B-C). These data strongly support the conclusion that APOE indeed coats the cytoplasmic-facing monolayer surface of LDs.

### APOE moves onto cytoplasmic LDs via ER membrane bridges

We hypothesized two possible routes by which APOE could traffic to the LD surface. In the first route, APOE first fully translocates into the ER lumen. It is then exported from the ER lumen to the cytoplasmic compartment via retrotranslocation to reach LDs. It was previously suggested that APOB retrotranslocates from the ER to the surface of LDs at ER-LD contact sites and is subsequently degraded (Ohsaki et al., 2006; Suzuki et al., 2012). In the other possible route, APOE avoids translocation into the ER, instead re-routing onto the cytoplasmic surface of the ER before targeting LDs at ER-LD contact sites. This second possibility is similar to the trafficking of membrane proteins such as DGAT2 that traffic to LDs via ER membrane bridges using the ERTOLD pathway (Wilfling et al., 2013; Song et al., 2020).

If APOE translocates into the ER before trafficking to LDs, we expect the APOE signal peptide to be cleaved. Therefore, we first examined whether the signal peptide of LD-associated APOE was cleaved by signal peptidase or retained. However, the signal peptide of APOE is quite small, with a molecular weight of approximately 1 kDa, and no corresponding mass shift of APOE was resolved via SDS-PAGE of OA treated cell lysates (Figure S1C). We instead employed an epitope-tagging strategy wherein a FLAG tag was appended to the N-terminal end of the signal peptide. Cells transfected with this construct and treated with OA had APOE signal that was positive for both FLAG and Emerald, indicating that the signal peptide was retained (Fig. S3A). By contrast, APOE in the secretory pathway was Emerald but not FLAG positive, indicating the signal peptide was cleaved in the luminal pool. This suggests that LD-targeted APOE has not been exposed to the ER lumen.

To test if APOE trafficking to LDs required retrotranslocation, we treated cells with the p97 inhibitor DBeQ concomitantly with OA treatment. No reduction in APOE signal on LDs was observed upon p97 inhibition (Fig. S3B). By contrast, inhibiting the translation of new protein during OA treatment with cycloheximide significantly attenuated the LD associated pool of APOE (Fig. S3C). This suggests that APOE routed to LDs is newly translated and not routed to LDs by p97-dependent retrotranslocation.

Since our results did not support trafficking of APOE to LDs after retrotranslocation from the ER, we next explored the second possibility: subversion of translocation into the ER followed by trafficking from the ER to LDs via membrane bridges. To track the targeting of APOE to LDs over time, we performed Airyscan live-cell imaging of APOE, LDs, and the ER during OA loading. We observed both half-circles and full rings of APOE forming around LDs at ER-LD contact sites (Fig. 3A; and Videos 1 and 2). Half-circles of APOE colocalized with ER wrapping around the surface of LDs. By contrast, only segments of APOE rings colocalized with the ER at a contact site, while other segments localized to the LD in the absence of ER. Our immunogold EM data also revealed ER-LD contact sites where gold labelling was observed both on the cytoplasmic face of the ER and on the surface of LDs (Fig. 3B). These data are consistent with a model in which APOE moves from the cytoplasmic face of the ER onto the surface of LDs via membrane bridges, as described for ERTOLD membrane proteins.

**Figure 3:**
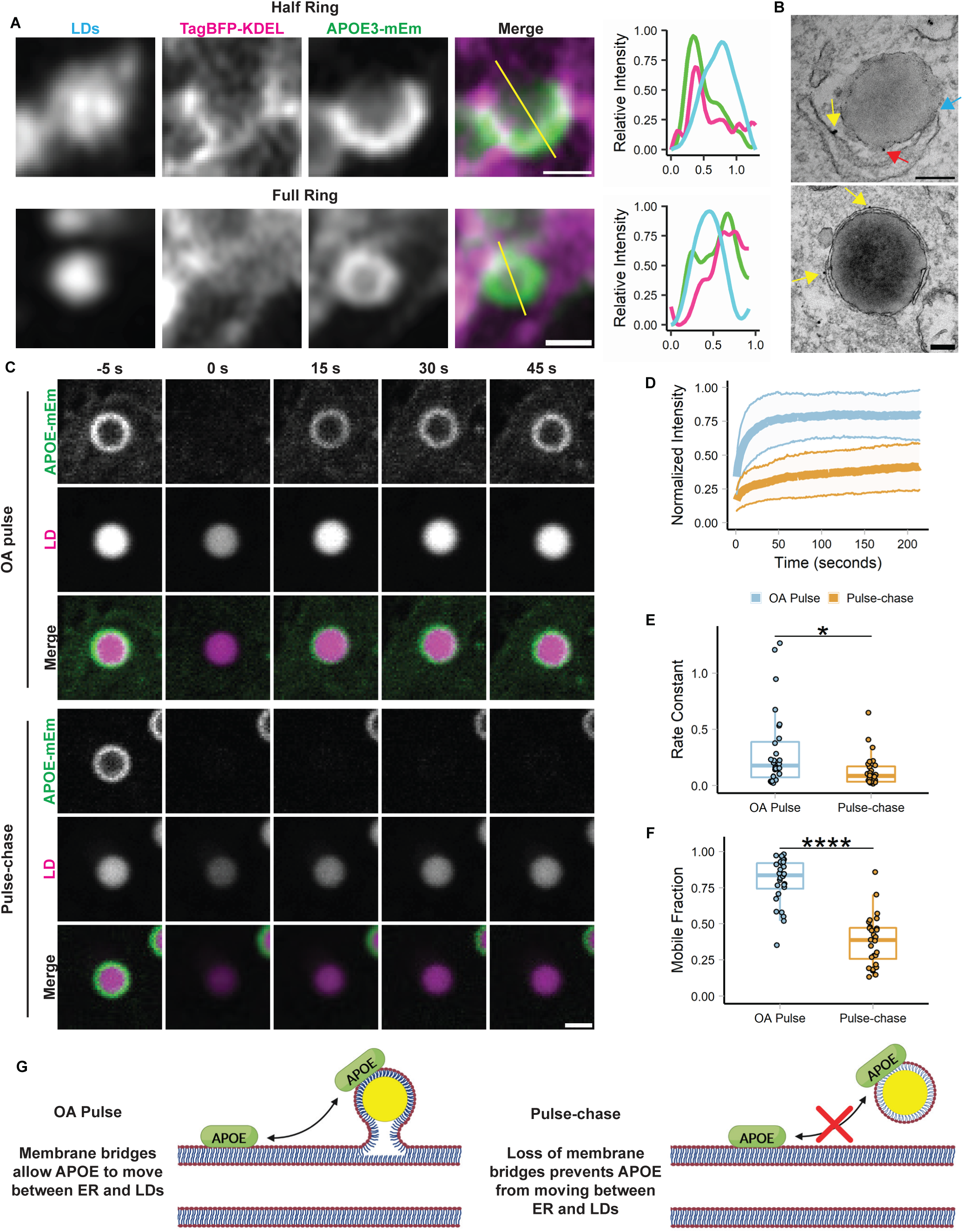
APOE targets LDs from the ER. (**A**) Representative frames from fast Airyscan movies showing the localization of LD-associated APOE relative to the ER after 4 hours of treatment with 400 µM OA in TRAE3-H cells. Cells were transfected with APOE3-mEm and the ER marker TagBFP2-KDEL and labelled for LDs with BODIPY 665/676. In the merged images, the ER is in magenta and APOE is in green. The yellow lines across the merged images indicate the line of pixels used to create the linescan graphs to the right of the images. In the linescan graphs, the relative fluorescence intensity of BODIPY-labelled LDs is in cyan, APOE3-mEm is green, and the ER is magenta. Two different localization patterns were observed: “half rings”, in which APOE partially covers the LD surface and colocalizes with the ER and “full rings”, where APOE fully encloses the surface of the LD and only partially colocalizes with the ER. Scale bars, 500 nm. **(B)** Immunogold electron micrographs of endogenous APOE localization at membrane contact sites between the ER and LDs in TRAE3-H cells treated with 400 µM OA for 5 hours. The blue arrow points to a direct membrane contact between the ER and an LD. Yellow arrows mark APOE localized to the cytoplasmic face of the ER membrane. The red arrow marks APOE localized to the cytoplasmic surface of the LD. Scale bars, 200 nm. **(C)** Representative frames from confocal FRAP movies of APOE3-Em on the surface of BODIPY 665/676 labelled LDs in primary rat cortical astrocytes during an OA pulse (200 µM OA for 4 hours) or an OA pulse-chase (200 µM OA for 4 hours followed by 2 hours chase in complete media – OA). APOE fluorescence was bleached at the 0 s timepoint. Scale bar, 1 µm. **(D)** Normalized intensity of APOE signal within the bleach ROI over time, with t = 0 sec denoting the time at which APOE was bleached. The bold center line is the mean normalized intensity, and the upper and lower bounds of the ribbon represent ± standard deviation (SD). N = 28 cells per condition, collected from three independent experiments. **(E)** Comparison of the rate constant of recovery *k* between OA pulse and pulse-chase conditions. The rate constant was derived by fitting each recovery curve to the equation y = C (1 – e^-^*^k^*^t^). N = 28 cells per condition, collected from three independent experiments. * p<0.05 **(F)** Comparison of the mobile fraction between OA pulse and pulse-chase conditions. The mobile fraction was derived by fitting each recovery curve to the equation y = C (1 – e^-^*^k^*^t^), where C is equal to the asymptote of the curve i.e. the mobile fraction. N = 28 cells per condition, collected from three independent experiments. **** p<0.0001. P-values were calculated via the Wilcoxon rank-sum test. **(G)** Schematic illustrating interpretation of the results of the FRAP experiment. When APOE on the LD is bleached during the OA pulse, it recovers very rapidly with a high mobile fraction. This indicates that bleached APOE on the LD is rapidly exchanged for unbleached APOE. After a short washout, LD-associated APOE recovers slowly or not at all, indicating that unbleached APOE molecules are unable to replace bleached ones on the LD. We hypothesize LD-APOE exchanges with APOE on the cytoplasmic face of the ER via membrane bridges during OA loading. These bridges are reduced or lost after OA washout, preventing exchange of APOE between LDs and the ER.

To further test our model, we performed fluorescence recovery after photobleaching (FRAP) experiments on LD-associated APOE. When APOE on LDs was bleached during loading with 200 µM OA, it rapidly recovered, nearly reaching its initial intensity. By contrast, LD-associated APOE recovery was profoundly attenuated after an OA pulse-chase in which cells were first loaded with 200 µM OA-supplemented media for 4 hours and then chased in unsupplemented complete media for 2 hours. (Fig. 3C-D; and Videos 3 and 4). During the OA pulse, APOE has both a significantly higher rate constant of recovery and mobile fraction (Fig. 3E-F). These observations suggest that during the period of OA-loading, bleached APOE molecules are rapidly replaced with unbleached molecules from the ER. Nascent LDs that form in response to OA treatment remain attached to the ER by membrane bridges immediately following biogenesis, which would allow APOE molecules on the ER to exchange with the LD pool. However, if OA is washed out in a pulse-chase experiment, then these nascent LDs mature and detach from the ER. Therefore, after the chase, unbleached molecules are unable to move onto LDs and exchange with the bleached ones. These data support a model in which APOE moves onto LDs via the ER at membrane bridges between the ER and LDs (Fig. 3G).

### APOE targeting to LDs requires its amphipathic C-terminal domain

To further dissect the mechanism of APOE targeting to LDs, we expressed different truncated variants of APOE and measured their capacity to target to the surface of LDs. APOE has two domains separated by a short hinge region. Its N-terminal domain forms a soluble 4-helix bundle and contains a patch of basic residues required for APOE binding to lipoprotein receptors. The C-terminal domain is an amphipathic alpha helix that mediates binding of APOE to the surface of lipoproteins (Westerlund and Weisgraber, 1993). To determine which domains are required for LD targeting, we made fluorescently tagged versions of both the N and C terminal domains (Fig. 4A). Additionally, we created versions of full-length APOE and the separate domains in which the N-terminal signal peptide was deleted (Δss). Deleting the signal peptide causes APOE to be only expressed in the cytoplasm. As we previously observed, full length APOE targeted LDs upon OA treatment. Full-length APOE Δss localized to the cytoplasm, and was enriched on the surface of LDs (Fig. 4B-C).

**Figure 4:**
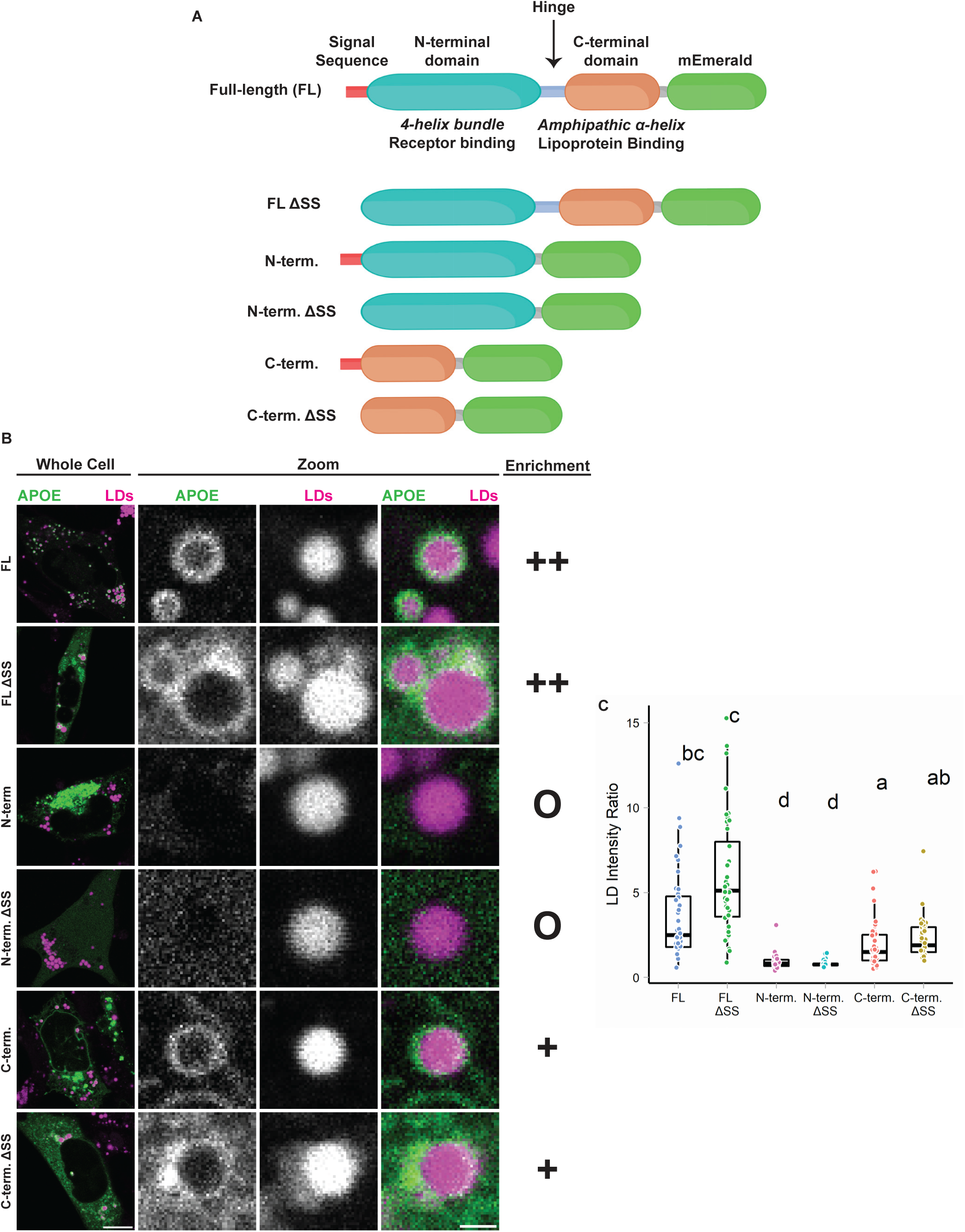
The C-terminal domain is required for LD targeting of APOE. (**A**) Schematic of the APOE truncation constructs used in this experiment. **(B)** Representative confocal slices of TRAE3-H cells transfected with the indicated construct, stained for LDs with BODIPY 665/676, and treated with 400 µM OA for 5 hours. “**O**” denotes no enrichment of signal on the LD surface, “**+**” indicates partial enrichment, and “**++**” indicates full enrichment. **(C)** Quantification of LD targeting of each construct. The LD Intensity Ratio was calculated by dividing the mean mEm fluorescence intensity on LDs divided by the mean mEm fluorescence intensity of the rest of the cell. Letters indicate significance groups. Conditions denoted by the same letter have no statistically significant difference. N = 40 cells per condition. Each data point represents one cell. Data was collected and pooled from three independent experiments. P-values were calculated via Dunn’s Test for pairwise multiple comparisons. FL, full-length APOE. N-term., N-terminal domain of APOE. C-term., C-terminal domain of APOE. ΔSS, construct has the N-terminal signal peptide deleted.

However, not every LD was coated with APOE, suggesting heterogeneity in binding affinity of APOE to LDs even when APOE is available for binding in the cytosol. The N-terminal domain with a signal sequence did not target LDs. The N-terminal domain Δss was soluble in the cytoplasm but did not exhibit LD surface enrichment (Fig. 4B-C). The C-terminal domain of APOE bound to LDs with or without a signal peptide, although its binding to LDs was less efficient than the full-length protein (Fig. 4B-C). We conclude that the C-terminal domain is necessary for binding of APOE to LDs, but inefficient for proper targeting as observed with the full-length protein.

### APOE regulates LD size

We next set out to identify the function of APOE on LDs. We hypothesized that APOE on LDs modulates LD metabolism. To test this, we performed OA pulse-chase assays in TRAE3-H cells transfected with an siRNA against *APOE* or a non-targeting control siRNA. A 5-hour OA pulse was used to stimulate LD biogenesis and drive APOE to LDs. OA-containing media was subsequently washed out and replaced with OA-free complete media for 18 hours, during which LDs were catabolized. BODIPY-stained LDs were imaged in live cells at the baseline, OA pulse, and pulse-chase timepoints. This design tests the effect of *APOE* knockdown on both the biogenesis and turnover of LDs.

At each time point, there was no difference in the area of LDs per cell, suggesting that *APOE* knockdown does not affect the total amount of neutral lipid in the cell. However, we observed a marked shift in the size distribution of LDs in *APOE* knockdown cells compared to control cells (Fig. 5A-D). After the OA pulse we observed larger and fewer LDs in APOE knockdown cells. This suggests that, although the same total amount of neutral lipid is present in these cells, LD biogenesis is altered, causing neutral lipids to concentrate in a smaller number of larger LDs. The altered size distribution is maintained after the pulse-chase.

**Figure 5:**
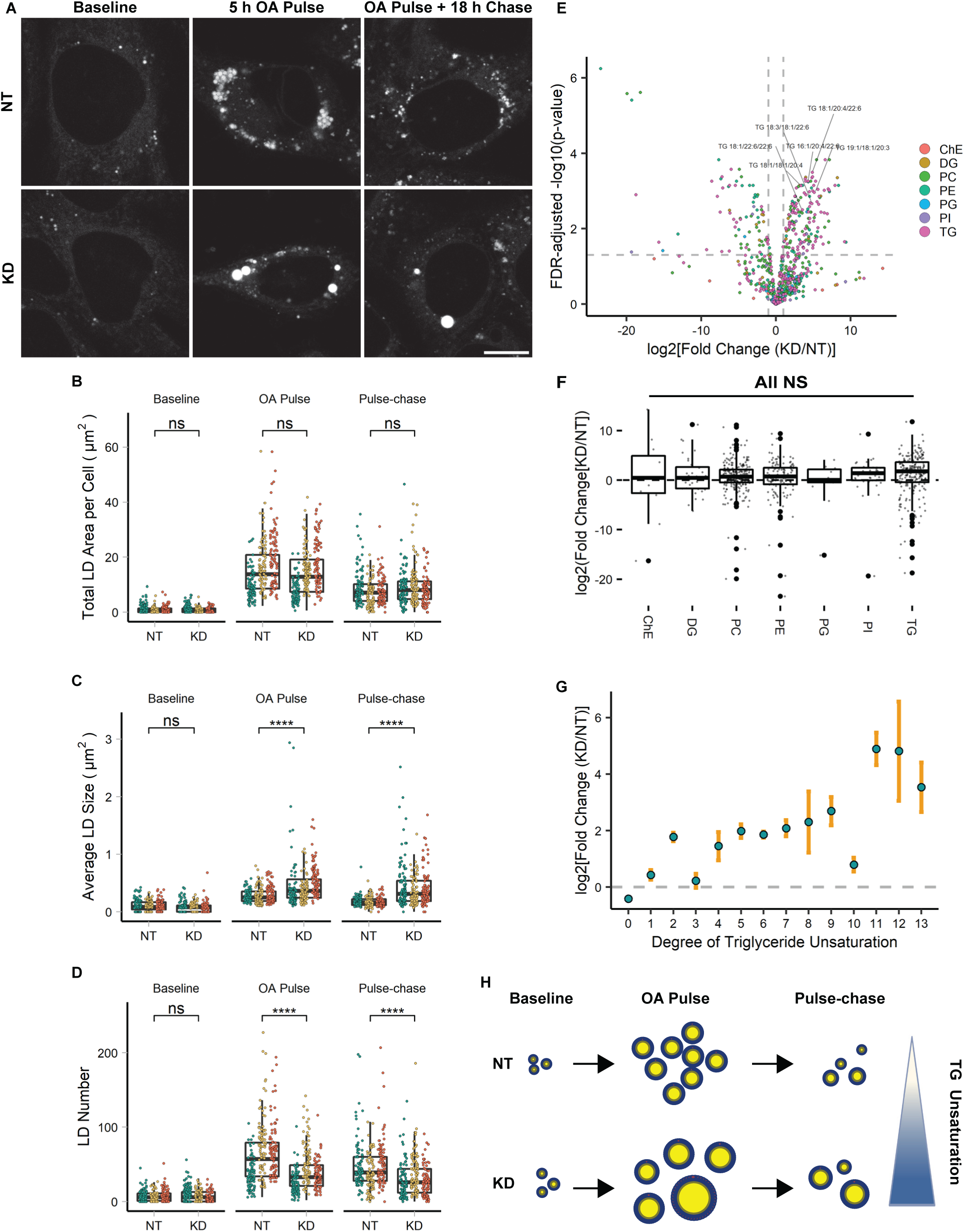
APOE modulates LD size distribution and triglyceride saturation. (**A**) Representative confocal slices of TRAE3-H transfected with non-targeting (NT) or *APOE* (KD) siRNA, stained for LDs with BODIPY 493/507, and imaged live at each timepoint of the OA-pulse chase assay. Scale bar, 10 µm. **(B-D)** Quantification of LD parameters in NT or *APOE* KD cells at each timepoint of the OA pulse-chase assay. (B) Total LD area was measured as the area of the entire LD mask per cell in µm^2^. (C) Average LD size was calculated as the mean LD area per cell in µm^2^. (D) Number of LDs per cell. Each data point represents one cell, and each color represents data collected from a separate, independent experiment. N = 90 cells per genotype, timepoint, and independent experiment. ns p>0.05, **** p<0.0001. P-values were calculated using a clustered Wilcox rank sum test via the Rosner-Glynn-Lee method and Bonferonni-corrected for multiple comparisons. **(E)** Volcano plot showing log_2_[Fold Change(KD/NT)] vs. –log_10_(p-value) of lipid species identified using untargeted whole cell lipidomics, comparing NT siRNA-transfected vs. *APOE* siRNA-transfected TRAE3-H cells after an OA pulse-chase. Lipids were measured by LC-MS/MS and normalized by both cell count and internal standards from 3 independent biological samples. P-values were calculated using limma-based differential enrichment analysis in the lipidR package. PC phosphatidyl choline, LPC lysophosphatidyl choline, LPE lysophosphatidyl ethanolamine, PE phosphatidyl ethanolamine, PI phosphatidylinositol, PS phosphatidylserine, DG diacylglycerol, TG triacylglycerol ChE cholesterol ester, DG diacylglycerol, PC phosphatidyl choline, PE phosphatidyl ethanolamine, PG phosphatidylglycerol, PI phosphatidylinositol, TG triacylglycerol. **(F)** Lipid set enrichment analysis (LSEA) of overall lipid classes enriched in *APOE* KD over NT cells was performed using the lipidR package in R. Lipids were grouped by class and ranked by log2[Fold Change(KD/NT)]. There was no significant overall up or downregulation of any major lipid class in *APOE* KD relative to NT cells after OA pulse-chase. **(G)** Relative enrichment of triglyceride species grouped by degrees of unsaturation (the total number of double bonds). Blue points represent mean log2[Fold Change(KD/NT)] and orange bars represent ± standard deviation. **(H)** Cartoon schematic illustrating results of **A-G.** In *APOE* KD cells, a smaller number of larger LDs form after OA pulse compared to NT cells. This LD size distribution phenotype is maintained after the pulse-chase, but there is no difference in total LD area. Moreover, *APOE* KD cells have more unsaturated triglyceride.

The distribution of LDs in fewer, larger LDs after an OA pulse suggests APOE knockdown causes defects in the formation and growth of LDs. One mechanism that affects LD size is their lipid composition. We hypothesized that APOE knockdown could impact the composition of LDs and have subsequent downstream effects on their size distribution. We performed whole cell lipidomics on TRAE3-H cells transfected with non-targeting siRNA or APOE siRNA and then subjected them to an OA pulse-chase. Numerous TG species were enriched upon APOE knockdown, many of which contained polyunsaturated fatty acids including arachidonic acid and docosahexaenoic acid (Fig. 5E). We observed no significant differences in the overall enrichment of TG, or any other major lipid class, in KD relative to control (Fig. 5F). This matches our previous microscopy data showing that total LD area is unchanged upon APOE depletion. However, comparing the relative abundance of TG species based on their degree of unsaturation, we found that APOE knockdown resulted in more TG species with higher degrees of unsaturation (Fig. 5G).

APOE knockdown alters the composition and size distribution of LDs formed during lipogenesis (Fig. 5H). However, global depletion of APOE via siRNA abrogates both LD-associated APOE and APOE in the secretory pathway, which is still observed in a subset of cells upon OA treatment (Fig. 1C). It is possible that the LD phenotypes we observed upon APOE knockdown are in part due to indirect effects of disrupting the balance of lipoprotein-based lipid secretion and uptake. To dissect the function of specifically LD-associated APOE, we performed rescue experiments with full-length APOE or a chimeric version of APOE that targets only to LDs. We made two rescue constructs, one in which synonymous mutations were introduced into the *APOE* sequence to impart RNAi resistance, and an “LD-only” chimeric construct in which the signal peptide of APOE was replaced with the LD-targeting hairpin domain of GPAT4 (Fig. 6A). The siRNA used in the knockdown studies targets the region of the *APOE* mRNA that encodes the signal peptide, and therefore removing the signal peptide makes this construct also resistant to knockdown. Both rescue constructs were tagged with HA at the C-terminus. The inclusion of a C-terminal tag prevented the APOE antibody from binding to its epitope, allowing us to distinguish between endogenous and exogenous HA-tagged APOE using anti-APOE or anti-HA antibodies (Fig. 6B). After APOE depletion, cells were transduced with lentivirus carrying either the full-length RNAi-resistant APOE or LD-only APOE and subjected to an OA pulse-chase. Western blot confirmed depletion of endogenous APOE and expression of both HA-tagged rescue constructs (Fig. 6B). Expression of either the RNAi-resistant full-length or the LD-only version of APOE fully rescued the LD size phenotype observed upon APOE knockdown (Fig. 6C-F). This supports a role for LD-associated APOE in modulating LD metabolism.

**Figure 6:**
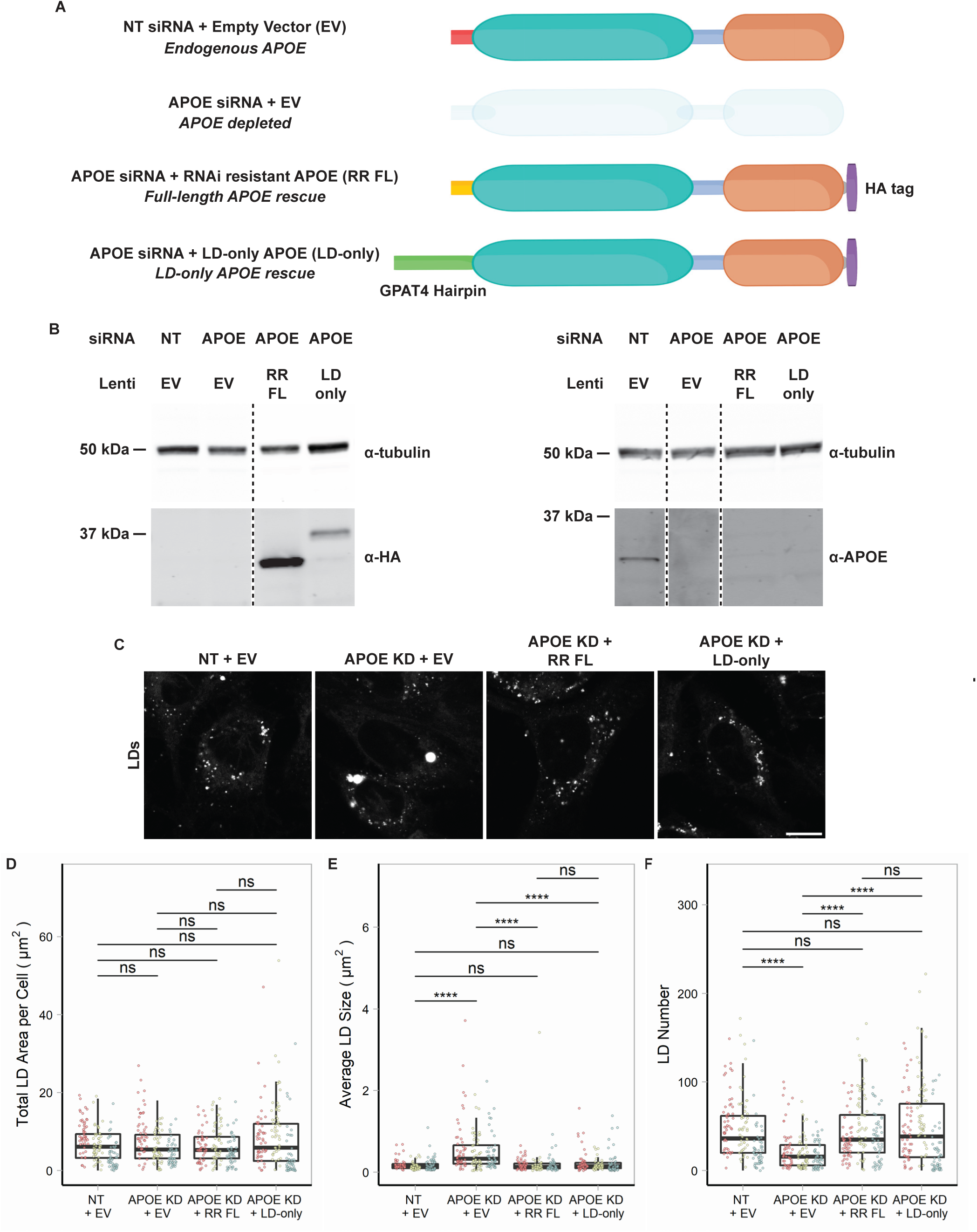
LD-associated APOE modulates LD size. (**A**) Cartoon illustrating the conditions used in the APOE rescue experiment. Endogenous APOE3 protein is present in cells transfected with a non-targeting siRNA. APOE protein is depleted upon *APOE* knockdown. The siRNA used to knock down *APOE* targets the mRNA sequence encoding the N-terminal signal peptide. The RNAi-resistant full-length APOE (RR FL) rescue construct consists of APOE3 with synonymous mutations in the signal peptide that impart resistance to the APOE siRNA. The LD-only APOE construct has the signal sequence removed, making it insensitive to the APOE *siRNA*, and replaced with the hairpin domain of the LD protein GPAT4. This version of APOE only targets LDs and never enters the ER lumen. **(B)** Western blot of lysates of TRAE3-H cells transfected with the indicated siRNA and transduced with the indicated lentivirus. The same samples were run on two separate SDS-PAGE gels, with 20 µg of total protein loaded into each well. Gels were transferred onto nitrocellulose membranes at the same time and then blotted with anti-HA or anti-APOE antibody together with an anti-tubulin antibody. Both the RR FL and LD-only APOE constructs are expressed in an endogenous APOE knockdown background. Moreover, the HA tag obstructs the epitope of the APOE antibody, allowing endogenous APOE and exogenous, HA-tagged APOE to be distinguished. **(C)** Representative confocal slices of cells transfected with non-targeting siRNA or *APOE* siRNA and transduced with an empty vector control, RR FL APOE, or LD-only APOE. Cells were subjected to an OA pulse-chase as described in **5A,** fixed, and stained for LDs with BODIPY 493/507. Scale bar, 10 µm. **(D-F)** Quantification of LD parameters the conditions described in (A) after an OA pulse-chase assay. (D) Total LD area was measured as the area of the entire LD mask per cell in µm^2^. (E) Average LD size was calculated as the mean LD area per cell in µm^2^. **(F)** Number of LDs per cell. Each data point represents one cell, and each color represents data collected from a separate, independent experiment. N = 60 cells per condition and independent experiment. ns p>0.05, **** p<0.0001. P-values were calculated using a clustered Wilcox rank sum test via the Rosner-Glynn-Lee method and Bonferonni-corrected for multiple comparisons.

### APOE4 expression results in large LDs with impaired turnover

We repeated the OA pulse-chase assay in targeted replacement astrocytes expressing human APOE3 (TRAE3-H) or APOE4 (TRAE4-H) to test the effect of APOE4 on LD biogenesis and turnover. APOE4 targeted LDs upon OA treatment to the same degree as APOE3, indicating that the E4 mutation does not compromise LD-targeting (Fig. S4). There were no significant differences in total LD area or LD number between TRAE3-H and TRAE4-H astrocytes at baseline, although E4 LDs were slightly larger. After OA pulse, E4 LDs were larger and fewer than E3 LDs, yet the total LD area remained the same, similar to the knockdown phenotype (Fig. 7A-D). However, after OA pulse-chase, E4 cells had a greater total LD area and larger LDs than E3, but no difference in LD number. This suggests that during LD biogenesis, neutral lipids partitioned into a smaller number of larger LDs in E4, similar to the phenotype we observed upon APOE knockdown. However, unlike the knockdown, the large LDs in *APOE4* cells displayed impaired turnover, as the total LD area was greater in *APOE4* cells after pulse-chase.

**Figure 7:**
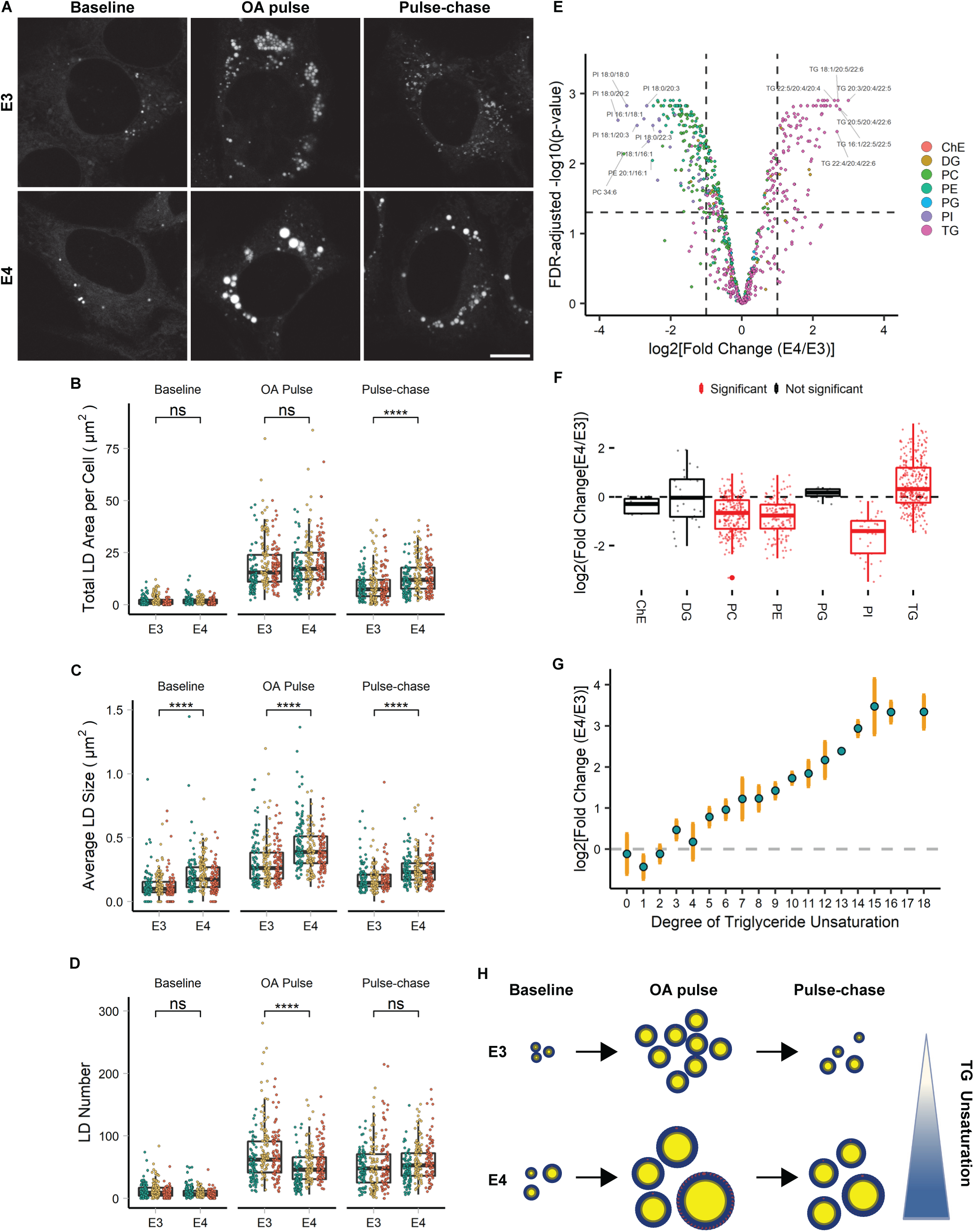
*APOE4* promotes large LDs with highly unsaturated triglyceride and impaired turnover. (**A**) Representative confocal slices of TRAE3-H or TRAE4-H cells labelled for LDs with BODIPY 493/507 and imaged live at each timepoint of the OA pulse-chase assay described in Figure 5A. Scale bar, 10 µm. **(B-D)** Quantification of LD parameters in TRAE3-H or TRAE4-H cells at each timepoint of the OA pulse-chase assay. (B) Total LD area was measured as the area of the entire LD mask per cell in µm^2^. (C) Average LD size was calculated as the mean LD area per cell in µm^2^. (D) Number of LDs per cell. Each data point represents one cell, and each color represents data collected from a separate, independent experiment. N = 90 cells per genotype, timepoint, and independent experiment. ns p>0.05, **** p<0.0001. P-values were calculated using a clustered Wilcox rank sum test via the Rosner-Glynn-Lee method and Bonferonni-corrected for multiple comparisons. **(E)** Volcano plot showing log_2_[Fold Change(E4/E3)] vs. –log_10_(p-value) of lipid species identified using untargeted whole cell lipidomics, comparing TRAE3-H vs. TRAE4-H after an OA pulse-chase. Lipids were measured by LC-MS/MS following normalization by cell count from 3 independent biological samples. P-values were calculated using differential enrichment analysis in the lipidR package. PC phosphatidyl choline, LPC lysophosphatidyl choline, LPE lysophosphatidyl ethanolamine, PE phosphatidyl ethanolamine, PI phosphatidylinositol, PS phosphatidylserine, DG diacylglycerol, TG triacylglycerol ChE cholesterol ester, DG diacylglycerol, PC phosphatidyl choline, PE phosphatidyl ethanolamine, PG phosphatidylglycerol, PI phosphatidylinositol, TG triacylglycerol. **(F)** Lipid set enrichment analysis (LSEA) of overall lipid classes enriched in E4 vs. E3 cells was performed using the lipidR package in R. Lipids were grouped by class and ranked by log2[Fold Change(E4/E3)]. Lipid classes colored red are significantly up or downregulated in E4. Classes colored black are not significantly different between E3 and E4. **(G)** Relative enrichment of triglyceride species grouped by degrees of unsaturation (the total number of double bonds). Blue points represent mean log2[Fold Change(E4/E3)] and orange bars represent ± standard deviation. **(H)** Cartoon schematic illustrating results of **A-G.** In *E4* cells, a smaller number of larger LDs compared to E3 form after OA pulse. After the pulse-chase, the large LDs observed in E4 have impaired turnover and are enriched in unsaturated triglyceride.

To measure how APOE4 expression alters cellular lipid metabolism, we performed whole-cell lipidomics on APOE3 and APOE4-expressing cells. We observed a significant increase in the abundance of TG in *APOE4* cells relative to *APOE3*, matching our microscopy data showing that E4 cells have more total LD area after OA pulse-chase (Fig. 7E-F). When examining individual lipid species, we observed an upregulation in various triglycerides in E4, while many phosphatidylcholine, phosphatidylethanolamine, and phosphatidylinositol species were less abundant in E4 (Fig. 7E). Much like we observed in *APOE* KD cells, many of the triglyceride species upregulated in E4 were highly unsaturated (Fig. 7G). We conclude that *E4* acts as a gain of toxic function allele with respect to its role on LDs, promoting the formation of large highly unsaturated LDs with impaired turnover (Fig. 7H).

### APOE4 LDs are more sensitive to lipid peroxidation

The physiological consequence of accumulating unsaturated TG in large LDs as observed in *APOE4* cells is unclear. Previous studies of LDs reconstituted *in vitro* demonstrated that having more unsaturated TG promotes larger droplets, similar to the phenotype we observed in cells (Lange et al., 2021). Moreover, these unsaturated LDs exhibit increased sensitivity to lipid peroxidation *in vitro*. We hypothesized that the enrichment of unsaturated TG in E4 LDs makes them more sensitive to lipid peroxidation. To test this hypothesis, we first performed the OA pulse-chase treatment in *APOE3* or *APOE4* cells as previously described. We then loaded cells with BODIPY C11—a fluorescent lipid peroxidation sensor (Fig. 8A). BODIPY C11 contains an 11-carbon fatty acid moiety that allows it to incorporate into lipid membranes (Drummen et al., 2002). BODIPY C11 shifts its fluorescence emission wavelength from red to green when oxidized in the presence of lipid peroxides, and the ratio of green to red BODIPY C11 fluorescence acts as a readout of lipid peroxidation in cell membranes. We found that after loading C11 into cells for 30 minutes followed by a 2-hour chase in Hank’s balanced salt solution (HBSS), the probe accumulated in LDs. The bright C11 signal of LDs was easily segmented from the rest of the cell, allowing for the measurement of peroxidation specifically in LDs. To test whether APOE4 increases sensitivity to lipid peroxidation, we subjected TRAE3-H or TRAE4-H cells to an OA pulse-chase, loaded LDs with BODIPY C11, and treated cells with vehicle (EtOH) or cumene hydroperoxide (CHP), a drug that causes lipid peroxidation. When treated with EtOH, both genotypes exhibited minimal lipid peroxidation with no significant difference between *E3* and *E4.* However, LDs in *APOE4* cells treated with CHP showed a much greater ratio of green to red fluorescence than LDs in CHP-treated *APOE3* cells (Fig. 8B-D). This suggests that the larger, unsaturated LDs of *APOE4* cells are more sensitive to lipid peroxidation than LDs in *APOE3* cells.

**Figure 8:**
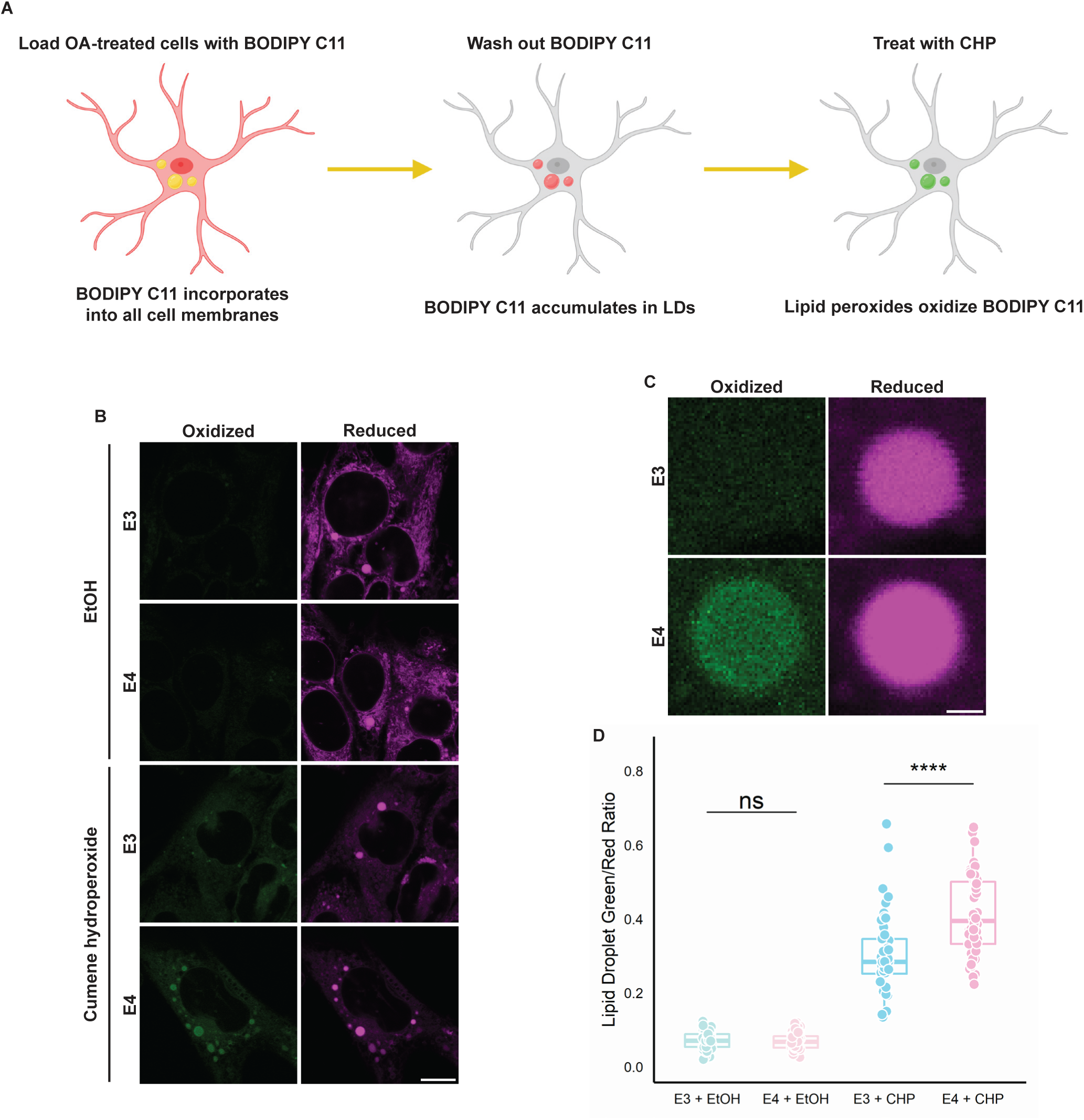
*APOE4* LDs are more sensitive to lipid peroxidation. (**A**) Schematic of a BODIPY C11-based assay for measuring the lipid peroxidation sensitivity of LDs. Cells are first subjected to an OA pulse-chase as described in Figure 5A, washed with HBSS, and then loaded with 2 µM BODIPY C11 in HBSS for 30 minutes. After 30 minutes, C11-containing HBSS is replaced with C11-free HBSS and cells are incubated for 2 hours, during which time C11 incorporates into LDs. Cells are then treated with 0.2% ethanol vehicle (EtOH) or 200 µM cumene hydroperoxide (CHP) for 2 hours and subsequently imaged. **(B)** Representative confocal slices showing TRAE3-H or TRAE4-H cells labelled with BODIPY C11 and treated with either 0.2% EtOH or 200 µM cumene hydroperoxide as described in **(A)**. The magenta channel shows reduced BODIPY C11 fluorescence, and the green channel shows the fluorescence of BODIPY C11 oxidized by lipid peroxides. Scale bar, 10 µm. **(C)** Inset of B showing difference in peroxidation of LDs between E3 and E4 cells. Scale bar, 1 µm. **(D)** Quantification of peroxidation in LDs in E3 or E4 cells treated with EtOH or CHP. The ratio was calculated by dividing the mean fluorescence intensity of green (oxidized) C11 fluorescence in the LD mask divided by the mean intensity of red (reduced) C11 fluorescence. ns p>0.05, **** p<0.0001. N = 50 cells per condition. Each data point represents one cell. Data was collected and pooled from three independent experiments. P-values were calculated via the Wilcoxon rank-sum test and Bonferonni-corrected for multiple comparisons.

## Discussion

Despite being the greatest genetic risk factor for Alzheimer’s disease, the detailed trafficking mechanisms of APOE in astrocytes have not previously been studied. Here, we show that APOE can divert from secretion and instead target LDs, where it modulates LD size and triglyceride saturation. LD-associated APOE coats the cytoplasmic-facing monolayer surface of LDs, and targets LDs from the cytoplasmic face of the ER membrane at ER-LD membrane bridges (Fig. 9A). LD-associated APOE is required during lipogenesis for maintaining LD size, while loss of APOE or APOE4 expression causes accumulation of polyunsaturated TG molecules within LDs.

**Figure 9:**
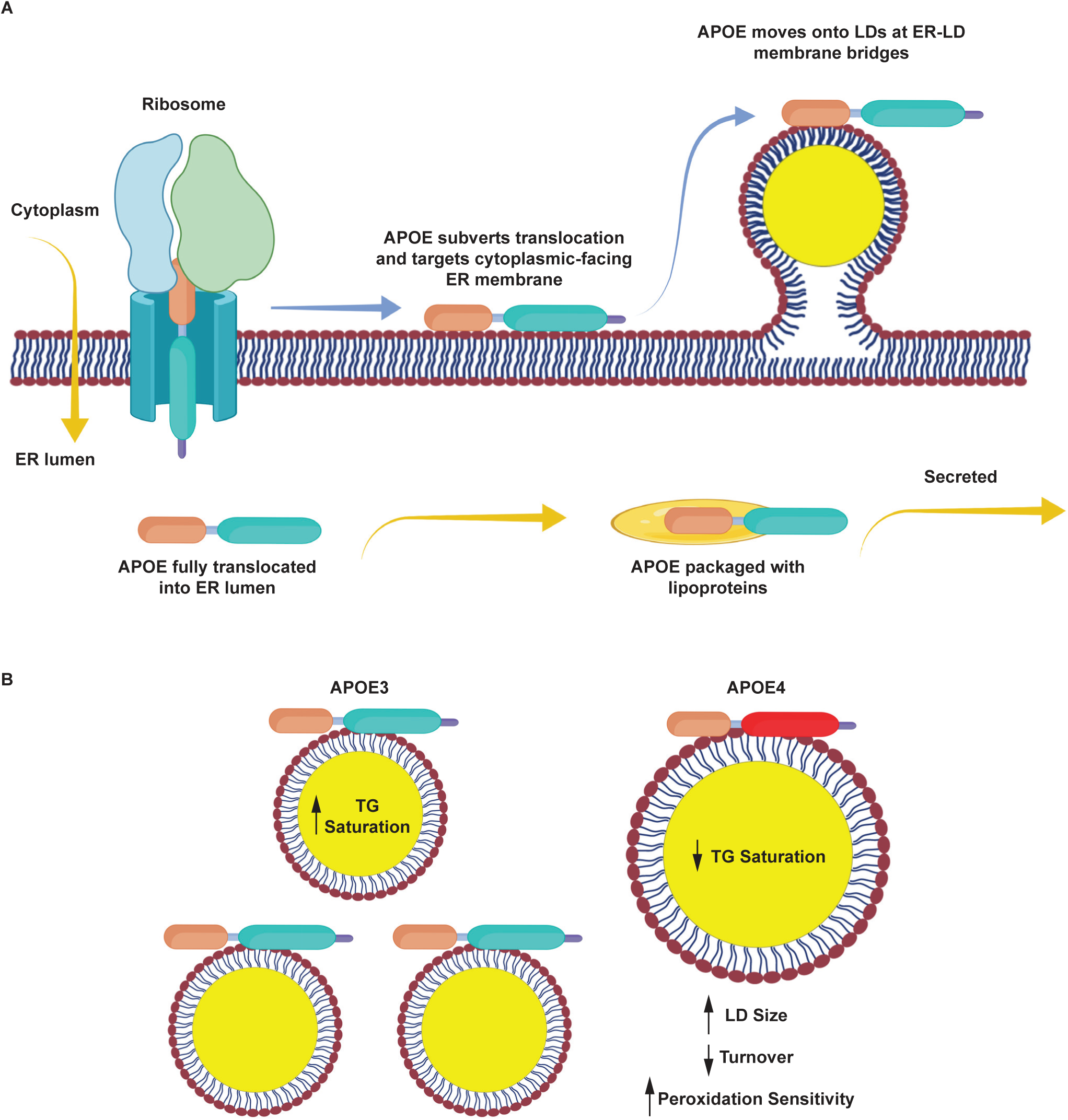
APOE targets cytoplasmic LDs via the ER and modulates LD composition and size. Schematic of our model of LD-associated APOE trafficking and function. **(A)** Normally, APOE is translocated into the lumen of the ER, where it assembles with nascent lipoprotein particles and is secreted. Under conditions that stimulate LD biogenesis, APOE subverts translocation via an unknown mechanism (retaining its signal peptide) and localizes to the cytoplasmic face of the ER membrane. It then moves onto LDs via membrane bridges between LDs and the ER. **(B)** On LDs, APOE is required to maintain triglyceride saturation and a dispersed LD size phenotype. In *APOE4* cells, LDs are larger, accumulate highly unsaturated triglycerides, have impaired turnover, and are more sensitive to lipid peroxidation. We hypothesize that defects in the function of LD-associated APOE4 promote lipid dishomeostasis and sensitize astrocytes to stress, which could facilitate the progression of Alzheimer’s pathology.

Importantly, the large unsaturated LDs in APOE4-expressing cells exhibit increased sensitivity to lipid peroxidation (Fig. 9B). The discovery that APOE traffics to LDs opens up many new avenues of investigation. What is the mechanism by which APOE is able to divert secretion and instead target LDs? How does LD-associated APOE regulate LD physiology? How do the LD defects observed in *APOE4* glial cells contribute to Alzheimer’s pathology?

The molecular mechanism by which APOE diverts from translocation into the ER lumen and instead targets LDs is unclear. Several other secreted proteins with lipid binding moieties can reroute to LDs. APOV and APOCIII have been shown to go to LDs (Shu et al., 2010; Gao et al., 2012; Sołtysik et al. 2019). One study demonstrated that the innate immune protein cathelicidin reroutes to LDs in response to OA loading or lipopolysaccharide treatment, retaining its N-terminal signal peptide (Bosch et al., 2020). Another apolipoprotein, APOB, possesses pause transfer sequences that cause it to stall during *in vitro* translocation, and parts of its nascent polypeptide chain becomes exposed to solution (Chuck et al., 1990; Kivlen et al., 1997). Pausing during translocation could act as a checkpoint for topological decision-making during translocation. The ER membrane complex (EMC) is involved in topological mediation of integral ER proteins (Hegde 2022). Proteins with uncleaved signal anchors can adopt orientations in which their N-terminus faces the cytoplasm or the ER lumen, a process regulated by EMC (Wu and Hegde 2023). EMC has also been implicated in topological decision making of ER hairpin proteins that target to LDs (Leznicki et al. 2021). Given the vast heterogeneity of targeting mechanisms during translocation, it is possible that some normally secreted proteins such as APOE utilize an uncharacterized conserved pathway to undertake alternative topological fates.

After avoiding translocation, APOE traffics to LDs from the cytoplasmic surface of the ER via membrane bridges. Recent study of the mechanism of protein trafficking from the ER to LDs identified both “early” and “late” ER to LD (ERTOLD) pathways (Song et al., 2022). Given that we observe APOE coating LDs several hours into OA treatment, we hypothesize that it may move onto LDs via the “late” pathway. However, unlike most ERTOLD proteins, LD binding of APOE is mediated by an amphipathic alpha helix rather than a hairpin domain. Deletion of the C-terminal amphipathic helical region of APOE ablates LD targeting. However, the C-terminus of APOE alone is less efficient at LD targeting than the full-length protein, suggesting a potential complementary role of the N-terminus. The N-terminal domain consists of four amphipathic alpha helices that form a bundle in solution (Chen et al. 2021). The N-terminal domain of APOE bound to the ER or LDs could retain this bundled fold or adopt an unfolded conformation, with the N-terminal amphipathic helices also mediating membrane association. This could explain the inefficient LD targeting observed in the C-terminal only construct.

Further study is necessary to uncover how LD-associated APOE controls triglyceride saturation and LD size. One possibility is that APOE itself stabilizes LDs and prevents their coalescence. The amphipathic alpha helices of APOE’s C-terminal domain could bind to monolayer packing defects and reduce surface tension. In the absence of APOE, higher surface tension would make lowering the surface area to volume ratio of LDs via Oswalt ripening more thermodynamically favorable (Thiam et al., 2013). The increase in unsaturated TG observed in *APOE* KD could be a downstream consequence of a change in LD size distribution, potentially due to impaired turnover of unsaturated TGs in larger droplets. Another possibility is that APOE acts as a scaffold, and the effects of APOE on LDs are due to a protein-protein interaction. The N-terminal domain of APOE forms a four helix-bundle that is homologous to the four-helix bundle domain of the perilipin family of LD proteins (Hickenbottom et al., 2004). In perilipins, this domain is involved in recruiting adipose triglyceride lipase to the surface of LDs. APOE has a patch of basic residues in helix 4 of its N-terminal domain that has previously been shown to mediate interactions between APOE and lipoprotein receptors. APOE may interact with other proteins on the surface of LDs, and this interaction could mediate the downstream effects of APOE on LD composition and size.

There is no consensus model regarding how *APOE4* increases an individual’s risk for developing Alzheimer’s disease. Prior work on APOE has focused on resolving how secreted APOE4 could lead to neurodegeneration (Martens et al. 2022). In this paper, we show that another pool of APOE exists on LDs that could also drive pathology. Further work is necessary to integrate the contributions of both pools of APOE to lipid homeostasis and Alzheimer’s disease. *APOE4* promotes the accumulation of unsaturated triglycerides in LDs. The large, unsaturated droplets in *APOE4* cells have impaired turnover and increased sensitivity to lipid peroxidation. Ischemia is correlated with both increased lipid peroxidation and neutral lipid accumulation in the brain (Yoshida et al., 1982; Gasparovic et al., 2001; Ioannou et al., 2019). Instances of ischemia are a major risk factor for AD, and increased sensitivity to hypoxic insult may play a role in driving pathology in *APOE4* (Elman-Shina and Efrati 2022). An accumulation of large LDs that are difficult to turnover and accumulate lipid peroxides could stimulate astrocytes to adopt a pro-inflammatory reactive state that precedes neurodegeneration. Lipid accumulation in other brain cell types has also been linked to neurodegeneration. LDs rich in cholesterol esters drive the accumulation of phosphorylated tau in iPSC-derived neurons (van der Kant et al., 2019). Aged microglia accumulate LDs and adopt a pro-inflammatory state (Marschallinger et al., 2020). The presence of these lipid-laden microglia correlates with an impaired response to stroke in aged mice (Arbaizar-Rovirosa et al. 2023). Although it remains to be determined whether APOE can traffic to LDs in microglia, we speculate that LD-associated APOE4 in microglia could also play a key role in AD pathology. This work adds to the body of evidence that is beginning to uncover key connections between APOE and intracellular glial lipid metabolism in driving Alzheimer’s disease.

## Materials and Methods

### Antibodies and chemicals

The following antibodies were used in this study. Dilutions used for western blot (WB), immunofluorescence (IF), and immunogold (IG) are given in parenthesis along with the manufacturer, catalog number, and RRID. Primary antibodies: recombinant rabbit anti-APOE mAb (1:300 IF; 1:50 IG; 1:500 WB; Abcam Cat# ab52607, RRID:AB_867704), mouse anti-HA-Tag mAb 6E2 (1:250 IF; 1:1000 WB; Cell Signaling Technology Cat# 2367, RRID:AB_10691311), mouse anti-DYKDDDDK Tag mAb 9A3 (1:200 IF; Cell Signaling Technology Cat# 8146, RRID:AB_10950495), mouse anti-GM130 mAb (1:250 IF; BD Biosciences Cat# 610822, RRID:AB_398141), mouse anti-Tubulin, alpha mAb DM1A (1:10,000 WB; Abcam Cat# ab7291, RRID:AB_2241126).

Secondary antibodies: donkey anti-rabbit IgG (H+L) Alexa Fluor 568 (1:500 IF; Thermo Fisher Scientific Cat# A10042, RRID:AB_2534017), donkey anti-mouse IgG (H+L) Alexa Fluor Plus 647 (1:500 IF; Thermo Fisher Scientific Cat# A32787, RRID:AB_2762830), donkey anti-rabbit IgG (H+L) IRDye 680RD (1:15,000 WB; LI-COR

Biosciences Cat# 926-68073, RRID:AB_10954442), donkey anti-mouse IgG (H+L) (H+L) IRDye 800CW (1:15,000 WB; LI-COR Biosciences Cat# 926-32212,

RRID:AB_621847), goat anti-rabbit IgG (H+L) Ultrasmall EM grade 0.8nm colloidal gold (1:100 IG; Aurion, Electron Microscopy Sciences Cat# 25101).

The following chemicals were used in this work: BODIPY 493/507 (ThermoFisher Cat# D3922), BODIPY 665/676 (ThermoFisher Cat# B3932), cycloheximide (Sigma Cat# C7698), DBeQ (Sigma Cat# SML0031), digitonin (Sigma Cat# D141), fibronectin (Sigma Cat# F4759), Image-iT™ Lipid Peroxidation Kit (ThermoFisher Cat# C10445), normal donkey serum (Sigma Cat# S30), 16% PFA solution, EM grade (EMS Cat# 15710), polybrene (Sigma Cat# TR-1003-G), poly-D lysine, 1.0 mg/mL (Sigma Cat# A-003-E), protease inhibitor cocktail (Sigma Cat# P8340), proteinase K (NEB Cat# P8107S), saponin (Fisher Cat# AAA1882014), sodium oleate (Sigma Cat# O7501).

### Plasmids

mEmerald-N1 (Addgene_53976) and mApple-SiT (Addgene_54948) were kind gifts from Dr. Michael Davidson (Florida State University). GFP-Plin2 was a kind gift from Dr. Alan Kimmel (described in Hsieh et al., 2012; National Institute of Diabetes and Digestive and Kidney Disease). The lentiviral vector pTK881 was a kind gift from Dr. Tal Kafri (UNC-Chapel Hill). TagBFP2-KDEL (Addgene_49150) was a kind gift from Dr. Gia Voeltz (University of Colorado, Boulder). LiveDrop-mEmerald was generated in our lab (Miner et al. 2023).

For plasmids constructed for this study, all PCRs were performed using Q5 High Fidelity DNA polymerase (M0419; New England Biolabs). A plasmid containing human APOE3-TurboGFP was purchased from Origene (Cat# RG200395). The APOE3 ORF was amplified from APOE3-TurboGFP and subcloned into an mEmerald-N1 backbone via Gibson assembly using HiFi DNA Assembly Master mix (New England Biolabs, E2621) to make APOE3-mEm. Only APOE3-mEm, not APOE3-TurboGFP was used for experiments in this study. APOE truncation mutants (APOE3-mEm ΔSS, APOE3-N-Em (1-191), APOE3-N-mEm ΔSS (19-191), APOE3-C-mEm, APOE3-C-mEm ΔSS were created from APOE3-mEm using Q5 Site-Directed Mutagenesis Kit (New England Biolabs, E0554S). The N-terminal domain constructs consist of residues 19-209 with or without residues 1-18 of the N-terminal signal peptide. C-terminal domain constructs consist of residues 234-317 with or without residues 1-18 of the N-terminal signal peptide. FLAG-ss-APOE3-mEm was created by adding an N-terminal FLAG tag to APOE3-mEm using Q5 Site-Directed Mutagenesis. The web tool “Synonymous Mutation Generator” was used to determine an optimal RNAi-resistant APOE sequence (Ong 2021). Synonymous mutations were then introduced into APOE3-mEm using Q5 Site-Directed Mutagenesis Kit. For lentiviral production, the RNAi-resistant APOE3 ORF was subcloned into pTK881 using Gibson assembly with a C-terminal HA tag added with PCR. HP-APOE3-mEm was created by amplifying the hairpin domain of GPAT4 (152-208) from LiveDrop-mEm and adding it to the N-terminus of APOE3-mEm ΔSS via Gibson Assembly. For lentiviral production, HP-APOE3 ORF was subcloned into pTK881 via Gibson Assembly with a C-terminal HA tag added with PCR.

### Cell culture and transfection

Immortalized targeted replacement astrocyte (TRAE3-H and TRAE4-H) cell lines were a kind gift from Dr. Patrick Sullivan of Duke University (described in Morikawa et al. 2005). U-2 OS cells were obtained from the UNC Tissue Culture facility. U-2 OS and targeted replacement astrocytes were maintained at 37°C and 5% CO_2_ in complete medium (CM) consisting of Dulbecco’s modified Eagle medium (DMEM) high glucose (Corning Cat# 15-013-CV), supplemented with 10% fetal bovine serum (FBS, VWR Cat# 97068-085), 2 mM glutamine (Corning Cat# 25005-CI), and 1X penicillin/streptomycin (Corning Cat# 30-002-CI). CM used for targeted replacement astrocytes was additionally supplemented with 200 µg/mL Geneticin (Gibco Cat# 10131027). Cells were confirmed free of mycoplasma and monitored at regular intervals using a Mycoplasma PCR Detection Kit (ABM G238).

Primary rat cortical astrocyte cultures were prepared from neonatal (1-2 day old) Sprague-Dawley rats (Charles River) as previously described (McCarthy and De Vellis 1980; Stogsdill et al., 2017). Briefly, cortices from both sexes were microdissected and digested with papain (10 units/brain, Worthington Biochemica Cat# LK003176l) for 25 min at 37°C. Tissue was washed three times with astrocyte growth media (AGM): DMEM high glucose (Corning Cat# 15-013-CV0), 10% FBS (VWR Cat# 97068-085), 2 mM L-glutamine (Corning Cat# 25005-CI), 1X penicillin/streptomycin (Corning Cat# 30-002-CI), 4 µg/mL hydrocortisone (Cayman Chemical Cat# 20739), 5 µg/mL bovine insulin (Sigma-Aldrich Cat# I6634), and 5 µg/mL N-acetyl-L-cysteine (Cayman Chemical Cat# 20261) and gently aspirated to remove residual papain. The tissue was triturated in AGM with three fire-polished Pasteur pipettes with progressively narrower bores and filtered through a 70-micron cell strainer (Falcon). Equal volumes of the cell suspension were plated on 75 cm^2^ flasks (Flacon) coated with 10 µg/mL of poly-D-lysine. Cells were plated such that 1 set of cortices was plated per flask. Flasks were maintained at 37°C, 5% CO_2_ with complete media exchanges on DIV1 and DIV2. On DIV3, confluent flasks were smacked by hand three times in pre-warmed PBS until the adherent astrocyte monolayer remained, and the flasks were replenished with AGM containing 10 µM AraC to limit proliferation of contaminating cells. On DIV6, flasks were replenished with fresh medium and subcultured on DIV7.

For light microscopy, cells were seeded in 8-well chamber slides with #1.5 high performance cover glass bottom (Cellvis C8-1.5H-N). 200 µL of filter-sterilized 10 µg/mL or 50 µg/mL poly-D-lysine was added to each well of an 8-well chamber slide for seeding primary astrocytes or targeted replacement astrocytes respectively. Slides were incubated at 37°C for 30-45 minutes and rinsed 3 times with milliQ water before seeding cells. For seeding U-2 OS cells, 200 µL of 10 µg/mL fibronectin was added to each well and slides were incubated at 37°C for 5 minutes and rinsed 3 times with milliQ water before seeding. The following cell numbers were used for seeding in 8-well slides: TRAE3-H/TRAE4-H, 12,000 cells per well; U-2 OS, 15,000 cells per well; primary astrocytes, 25,000 cells per well.

Plasmid DNA was transfected into all cell types using Lipofectamine 2000 (Invitrogen Cat# 11668019) according to the manufacturer’s instructions. A 2:1 ratio of Lipofectamine to plasmid DNA was used for all transfections. siRNAs were transfected using DharmaFECT 1 (Horizon Discovery Cat# T-2001) according to the manufacturer’s instructions. 5 µM siRNA was used for each transfection. Non-targeting (NT) siRNA (Cat# D-001810-04), APOE siRNA #1 (Cat# D-006470-02), and APOE siRNA #2 (Cat# D-006470-04) were purchased from Horizon Discovery. After transfection, cells were incubated in their corresponding “imaging media” lacking phenol red and antibiotics.

### Immunofluorescence sample preparation

Cells were washed twice in 1x PHEM buffer (60 mM PIPES, 27.3 mM HEPES, 8.22 mM MgSO_4_, 10 mM EGTA, pH 7.0) and fixed for 10 minutes at room temperature in room temperature-equilibrated 4% PFA in 1x PHEM. After fixation, cells were washed three times in 1x PHEM and then permeabilized in 0.01% saponin in PHEM for 10 minutes at room temperature. Permeabilization buffer was then replaced with blocking buffer (10% Normal Donkey Serum, 3% bovine serum albumin (BSA), 300 mM glycine, 0.01% saponin in PHEM) in which cells were incubated 30-45 minutes at room temperature.

Blocking solution was removed and cells were washed twice with 0.01% saponin in PHEM before applying primary antibody solution. Primary antibodies were diluted in antibody dilution solution (3% BSA, 0.01% saponin in PHEM). Cells were incubated in primary antibody solution for 24-48 hours at 4 °C. After primary antibody incubation, cells were washed with 0.01% saponin three times for 10 minutes each at room temperature. Secondary antibodies were diluted in antibody dilution solution together with 1 µg/mL BODIPY 493/507 to stain for LDs. Cells were incubated in secondary antibody solution for 1 hour at room temperature, then washed 3 times in PHEM for 10 minutes each at room temperature. Cells were imaged immediately after preparing or stored at 4 °C and imaged within a week of preparation.

### Immuno-gold electron microscopy

Targeted replacement cells were grown on Nunc Lab-Tek Permanox chamber slides. Cells were treated with 400 µM OA for 5 hours and fixed for 1 hour at room temperature in 4% paraformaldehyde in 0.15 M sodium phosphate buffer (PB). After three washes in 0.15 M PB the cells were immunostained using a modified protocol for pre-embedding immunogold-silver staining developed by Yi et al. (2001). Cell permeabilization and free-aldehyde inactivation was achieved using 0.1% saponin in 0.2 M glycine in 0.15 M PB, for 30 minutes, following by a 1-hour incubation in an Aurion Goat Blocking Solution (Aurion, Electron Microscopy Sciences, Fort Washington, PA). The cells were incubated in anti-APOE antibody overnight at 4°C (Rabbit anti-APOE, abcam ab52607 diluted 1:50 with IEM buffer (PB with 0.2% BSA-Ac/0.01% saponin). After washes in IEM buffer, the cells were incubated overnight at 4°C in a 1:100 dilution of Goat anti-rabbit IgG (H+L) Ultrasmall EM grade 0.8 nm colloidal gold (Aurion, Electron Microscopy Sciences, Fort Washington, PA, cat #25121-0.6ml). Following washes in IEM buffer, the cells were then washed with PB, post fixed in 2% glutaraldehyde in 0.15 M PB and held at 4 degrees for several days. Cells were rinsed in 0.15 M PB, and silver enhanced for 90 minutes using an Aurion R-Gent SE-EM Silver Enhancement Kit (Aurion, Electron Microscopy Sciences, Fort Washington, PA, cat #25521), followed by post-fixation in 0.1% osmium tetroxide for 15 minutes, gradient dehydration in ethanol and embedment in Polybed 812 epoxy resin (Polysciences, Inc., Warrington, PA). Grid-mounted 80 nm ultrathin sections were stained with 4% uranyl acetate and examined by transmission electron microscopy using a JEOL JEM-1230 transmission electron microscope operating at 80 kV (JEOL USA, Peabody, MA) equipped with a Gatan Orius SC1000 CCD camera and Gatan Microscopy Suite 3.0 software (Gatan, Inc., Pleasanton, CA).

### Western blot

Cells were lysed on ice in RIPA lysis buffer (50 mM Tris-HCl, 150 mM NaCl, 1% Triton X-100, 0.5% sodium deoxycholate, 0.1% SDS) supplemented with Protease Inhibitor Cocktail (Sigma) for 10 minutes. Adherent samples were scraped, collected, and centrifuged at 17,000xg for 10 minutes at 4 °C to isolate the post-nuclear supernatant. Protein concentrations in samples were measured using a Detergent Resistant Bradford Assay (ThermoFisher). Samples were prepared for SDS-PAGE in 6x Laemmli buffer and denatured for 5 minutes at 95 °C. Equal protein amounts were loaded into each well of a NuPAGE™ 10%, Bis-Tris, 1.0 mm, Mini Protein Gel (ThermoFisher). After SDS-PAGE, proteins were wet transferred to nitrocellulose for 1 hour at 100 V. Membranes were blocked in 5% milk in TBS for 30 minutes at room temperature and then incubated with primary antibodies diluted in 3% BSA in TBST overnight at 4 °C. Membranes were then washed 3 times with TBST and then incubated with secondary antibody solution for 1 hour at room temperature, washed 3x with TBST and imaged using an Odyssey CLx (LI-COR Biosciences).

### ELISA

Secreted APOE concentrations were determined using a human APOE ELISA kit (Abcam) per the manufacturer’s instructions. Briefly, 50 µL of 1 mL neat culture media was assayed and absorbance values were collected at 450 nm on a Synergy HT plate reader (BioTek).

### Fluorescence protease protection assay

Fluorescence protease protection assay was performed using a modified version of the protocol outlined in (Lorenz et al., 2006). Primary rat astrocytes were seeded on DIV7 and transfected with TagBFP2-KDEL and either mEm N1, APOE3-mEm, GFP-PLIN2, or LiveDrop-mEm. After transfection, media was replaced with imaging media supplemented with 50 ng/mL BODIPY 665/67. 24 hours post-transfection, cells were washed twice with KHMN (110 mM potassium acetate, 20 mM HEPES, 2 mM MgCl_2,_ 10 mM NaCl) buffer and placed in 150 µL KHMN. For each well, a field of view containing 2-3 cell transfected with TagBFP2-KDEL and mEm or the LD protein marker of interest was selected and a baseline image was taken. Optimization experiments determined 30 µM digitonin was the lowest concentration that showed full loss of cytoplasmic mEmerald signal in most cells after one minute, and this concentration was used in subsequent experiments. For each experiment, 150 µL of 60 µM digitonin in KHMN buffer was added to 150 µL of KHMN in the well for a final concentration of 30 µM digitonin and cells were allowed to permeabilize for one minute. Cells were re-focused and the “post-permeabilization” image was taken. 300 µL of 100 µg/mL proteinase K in KHMN was then added to the well for a final concentration of 50 µg/mL proteinase K. After 1 minute, cells were refocused and the “post-PK” timepoint image was taken. 3-5 wells per condition were performed per independent experiment, and data was collected and pooled form three independent experiments.

### FRAP

FRAP experiments were conducted in primary rat cortical astrocytes transfected with APOE3-mEm and stained for LDs with BODIPY 665/676 as described above.

Experiments were performed after treating cells for 4 hours in antibiotic and phenol red-free AGM supplemented with 200 µM OA for 4 hours (OA pulse) or cells treated with OA for 4 hours followed by a 2-hour chase in OA-free AGM. In each experiment, bleaching was conducted in 8-10 cells per condition, and only one APOE ring was bleached and analyzed per cell. Images were acquired once per second. Imaging started 5 seconds before bleaching and continued until 5 minutes post-bleach.

### OA pulse-chase assays

For APOE knockdown OA pulse-chase assays, 6,000 TRAE3-H cells were seeded in each well. 24 hours after seeding, cells were transfected with 5 µM of the indicated siRNA with Dharmafect 1 using the manufacturer’s protocol in imaging media. 48 hours post-transfection, media was replaced with imaging media supplemented with 200 ng/mL BODIPY 493/507 (for baseline wells) or imaging media supplemented with 400 µM sodium oleate (for +OA and washout wells) and 200 ng/mL BODIPY 493/507. After 5 hours of OA treatment, pulse-chase wells were replaced with imaging media, and “baseline” and “+OA” wells were imaged immediately.

For experiments comparing TRAE3-H and TRAE4-H cells, 12,000 cells were seeded in each well. 48 hours after seeding, the OA pulse-chase assay was conducted as described above.

For rescue experiments, 6000 TRAE3-H cells were seeded in each well. 24 hours after seeding, cells were transfected with siRNA as indicated above. 24 hours after siRNA transfection, cells were treated with the indicated lentivirus (UNC Lenti-shRNA Core Facility) diluted 1:10 in imaging media supplemented with 10 µg/mL polybrene. 48 hours after transduction, the OA pulse-chase was performed, with cells prepared for immunofluorescence at each timepoint.

### Lipid peroxidation assays

Cells first underwent the OA pulse-chase treatment as described above. 18-hours after the CM chase, cells were washed once in HBSS (ThermoFisher Cat# 14025092) and then loaded with 2 µM BODIPY C11 for 30 minutes. After loading, cells were washed once with HBSS and incubated in HBSS for 2 hours. Cells were then treated with 200 µM cumene hydroperoxide or 0.2% EtOH vehicle control for 2 hours and immediately imaged live.

### Light microscopy image acquisition

Confocal and Airyscan images were acquired using an inverted Zeiss 800/Airyscan single point scanning confocal microscope equipped with 405 nm, 488 nm, 561 nm, and 647 nm diode lasers, two Gallium Arsenide Phosphide (GaAsp) detectors and one Airyscan detector. Images were acquired using a Plan-Apo 63x/1.4 NA oil objective lens using ZEN Blue software. All live cell imaging was conducted at 37 °C and 5% CO_2_.

### Light microscopy image analysis

LD number and size analysis was performed using a semi-automated pipeline in Fiji (<u>Schindelin</u> et al., 2012). Individual cells were first manually segmented and the area outside of each cell was removed using the “Clear Outside” function. To segment LDs, the BODIPY channel was first passed through Gaussian filter followed by a Laplacian of Gaussian (LoG) filter. The radius of the LoG filter and the Gaussian blur was heuristically optimized for each image to maximize segmentation accuracy. Auto-thresholding using the Otsu algorithm was then applied, followed by the binary operations “Open”, “Fill Holes”, and “Watershed”. Then, “Analyze Particles” was used to measure the number and size of segmented LDs, as well as the sum area of all LDs in the cell.

For colocalization analysis, the APOE was segmented by first applying a median filter with a radius of 3 pixels to APOE channel followed by auto thresholding using the “Moments” algorithm. The Golgi was masked by applying a median filter with a radius of 2 pixels to the GM130 channel followed by auto thresholding using the “Huang2” algorithm. The Image Calculator function was used to create a mask of pixels were APOE and GM130 masks overlap. The Mander’s coefficient was calculated by dividing the area of the overlap mask by the area of the APOE mask.

To measure LD protein enrichment, the above algorithm was first applied to create an accurate mask of LDs in the cell. Then, the LD mask was expanded twice using the “Dilate Function” to include protein surrounding the surface of the LD. A second mask was created by subtracting the expanded LD mask from a mask of the entire cell. The mean intensity of the LD protein of interest was measured in the expanded LD mask as the subtracted whole cell mask. LD ratios were calculated by dividing the mean intensity of the LD mask by the mean intensity of the Cell-LD mask.

FRAP experiments were analyzed as described in (Day et al., 2012). The mean fluorescence intensity of the entire cell, the bleached ROI, and the background outside the cell was measured for each frame. The normalized intensity at frame *t* was calculated using the equation:

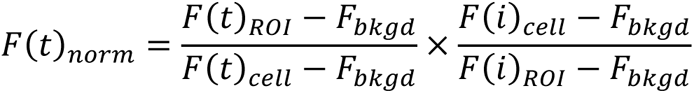

The normalized ROI intensity for each movie was fit to the equation:

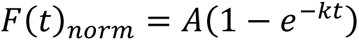

and the coefficients A (the mobile fraction) and k (the rate constant) were extracted from each.

### Lipidomics

Targeted replacement astrocytes were seeded in 6 cm dishes; 3 plates were used per condition. Cells were subjected to OA pulse-chase, washed 3 times with ice-cold PBS, scraped, and collected into 1.5 mL Eppendorf tubes. Cells were pipetted up and down 80-100 times on ice and counted using a Countess II automated cell counter (ThermoFisher) for normalization; 4 counts per plate were collected and averaged to ensure accuracy. Cells were then pelleted at 1000xG for 10 minutes and flash-frozen in liquid nitrogen. Cell pellets were extracted via a liquid-liquid partition with water (200 µL), methanol (300 µL), and methyl-tertbutyl ether (1 mL). An internal standard, Equisplash by Avanti Polar Lipids (Alabaster, AL), was spiked into the methanol used for extraction at 1.5 µg/mL. Samples were shaken for 15 minutes and then centrifuged at 20,000 rcf for 10 minutes. The top layer was dried and reconstituted with 150 µL of isopropanol for analysis.

Lipidomics analysis was performed using a Q Exactive HF-X (ThermoFisher, Bremen, Germany) coupled to a Waters (Milford, MA) Acquity H-Class UPLC. A 100 mm x 2.1 mm, 2.1 µm Waters BEH C18 column was used with the following mobile phases: A-60/40 ACN/H20 B-90/10 IPA/ACN; both mobile phases had 10 mM Ammonium Formate and 0.1% Formic Acid. A flow rate of 0.2 mL/min with a gradient starting at 32% B, which increased to 40% B at 1 min (held until 1.5 min) then 45% B at 4 minutes. This was increased to 50% B at 5 min, 60% B at 8 min, 70% B at 11 min, and 80% B at 14 min (held for 2 min). At 16 min the composition switched back to starting conditions and was held for 4 min. Samples were analyzed in positive/negative switching ionization mode with top 5 data-dependent fragmentation utilizing a stepped collision energy of 25, 35, and 45 V. A resolution of 60,000 was utilized with a scan range of 200–1200 m/z. Data-dependent acquisition was acquired at a resolution of 15,000 and an isolation window of 1.5 *m/z*.

### Lipidomics data processing

LC-MS data were analyzed by LipidSearch 4.2 by ThermoFisher Scientific and the peak area was normalized to the area of deuterated internal standards from the Avanti Equisplash mix as well as the number of cells. Lipids were identified by MS2 fragmentation with the following parameters: precursor tolerance 5 ppm, product tolerance 8 ppm, m-score threshold 2.0, and product relative intensity threshold 0.1%. The higher-energy C-trap dissociation and labeled databases were used for identification. The identifications were generated individually for each sample and then aligned by grouping the samples. Data was normalized to both internal standards and cell counts for each sample.

### Statistical analysis

RStudio was used to perform all statistical analysis (RStudio Team 2022). All plots were created using ggplot2 (Wickham 2009). The Rosner-Glynn-Lee variation of the Wilcoxon rank-sum test for clustered data was applied to asses statistical significance between groups from data collected from multiple independent experiments performed at different times. This test treats data from each independent experiment as a “cluster” and accounts for variability both within and between independent experimental trials (Rosner et al., 2006). The test was applied using the “clusrank” package in R, and was used for LD enrichment in Figures 1 and S4, LD parameter analysis in Figures 5, 6, and 7, and colocalization analysis in Figure 1 (Jiang et al. 2020).

For LD enrichment analysis in Figure 2, data was pooled from 3 independent experiments and p-values were calculated using standard Wilcox rank sum test and Bonferonni-corrected. Comparisons of FRAP data in Figure 3 was conducted using Wilcox rank sum tests. For LD enrichment analysis in Figure 4, the Dunn test was used to calculate pairwise p-values, and significance groups were assigned based on these pairwise p-values. Lipid peroxidation data in Figure 9 was compared using Wilcox rank sum tests and Bonferonni-corrected for multiple comparisons.

For lipidomics experiments, analysis was conducted using the lipidR Bioconductor package (Mohamed et al., 2020). P-values for volcano plots were calculated using the differential enrichment function in the lipidR package, which is based on the *limma* package and uses a moderated t-statistic and corrects for multiple comparisons using the false discovery rate (FDR). Lipid set enrichment analysis was conducted using the “lsea” function, based on gene set enrichment analysis (GSEA), with lipid species ranked by their log2FC.

## Supplemental Material

Figs. S1 and S2 show supporting data for Fig. 1. Fig. S3 shows supporting data for Fig. 1. 3. Fig. S4 shows supporting data for Fig. 7. Video 1 and Video 2 show partial and full rings of APOE on LDs at ER-LD contact sites and correspond to Fig. 3. Video 3 and Video 4 show FRAP movies corresponding to Fig. 3. Table S1 contains data and statistical analysis for the APOE knockdown lipidomics experiment. Table S2 contains the results of lipid set enrichment analysis on the APOE knockdown lipidomics data. Table S3 contains data and statistical analysis for the APOE3 vs APOE4 lipidomics experiment. Table S4 contains the results of lipid set enrichment analysis on the APOE3 vs APOE4 lipidomics data.

## Acknowledgements

We thank Patrick Sullivan (Duke University) and David Holtzman (Washington University in St. Louis) for providing the targeted replacement astrocyte cell lines; Victoria Madden of the Microscopy Services Laboratory for assistance with immunogold EM experiments; the University of North Carolina Department of Chemistry Mass Spectrometry Core Laboratory for their assistance with mass spectrometry analysis; Wendy Salmon, director of the Hooker Imaging Core, for assistance with fast Airyscan imaging; Sunghyun Lee and Tal Kafri for lentivirus production; Gregory Miner, Maria Clara Zanellati, and current Cohen Lab members for helpful discussions and feedback on the manuscript.

Research reported in this publication was supported by the National Institute of General Medical Sciences of the National Institutes of Health under award numbers R35 GM133460 (S.C.) and 5T32GM119999 (I.W.), by the National Institute on Aging under award number F31 AG069419 (I.W.), and by the Alzheimer’s Association under award number 2018-AARG-590347. Mass spectrometry work conducted by the UNC Mass Spectrometry Core Facility was supported by the National Science Foundation under Grant No. (CHE-1726291).

The authors declare no competing financial interests.

## Author Contributions

I.A. Windham and S. Cohen conceptualized the project, designed the research, acquired funding, and wrote the manuscript. J.V. Ragusa performed ELISA experiments and prepared primary astrocyte cultures. E.D. Wallace performed lipid extraction, data processing and mass spectrometry analysis. C.H. Wagner contributed to APOE truncation experiments. K.K. White performed immunogold electron microscopy. I.A. Windham performed all other experiments in the manuscript. I.A. Windham performed image analysis and wrote scripts for image and statistical analysis. S.C. supervised the research.

---

**Supplementary Figure 1:**
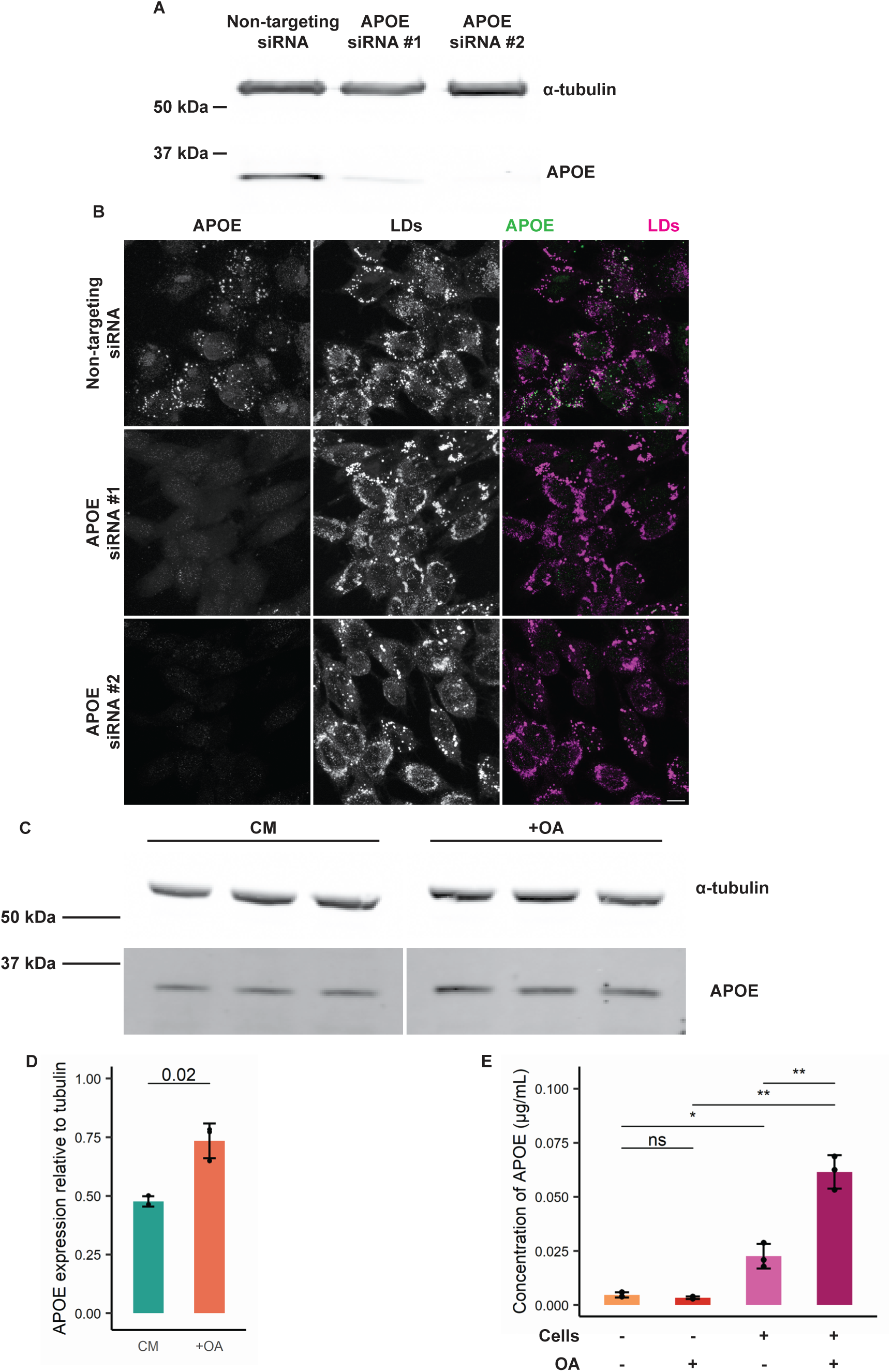
Validation of APOE antibody via siRNA-mediated knockdown and measurement of intracellular and secreted APOE ± OA. **(A)** Western blot of lysates of TRAE3-H cells transfected with a non-targeting siRNA or one of two different siRNAs against *APOE*. APOE siRNA #2 demonstrated the most thorough knockdown of APOE, and was used for subsequent loss of function studies in Main Figures 5 and 6. The antibody used to probe for endogenous APOE is the same used for both the immunofluorescence and immunogold experiments. **(B)** Representative confocal slices of TRAE3-H cells transfected with NT siRNA or one of two APOE siRNAs and treated with 400 µM oleate for 5 hours. Cells were fixed and stained for endogenous APOE with an anti-APOE antibody and for LDs with BODIPY 493/507. Little to no endogenous APOE signal was observed by immunofluorescence upon APOE knockdown. Scale bar, 10 µm. **(C-D)** (C) Western blot of lysates of TRAE3-H cells incubated in complete media (CM) or treated with 400 µM OA in CM for 5 hours (+OA). 10 µg of protein was loaded into each well. (D) APOE band intensity was quantified in ImageJ and normalized relative to alpha-tubulin band intensity. Intracellular protein levels of APOE significantly increase after OA treatment. N = 3 independent biological replicates. Data are expressed in bar graphs as means, and error bars represent ± standard deviation. P-value calculated using an unpaired, two-sample t-test. **(E)** TRAE3-h cells were treated with CM or CM supplemented with OA. Conditioned media was collected after 5 hours, and ELISA was performed to measure the concentration of secreted APOE. N = 3 independent experiments. For each experiment, samples were collected and measured in triplicate; each data point is the mean concentration of triplicate samples per experiment. Data are expressed in bar graphs as means, and error bars represent ± standard deviation. ns, p< 0.05, * p < 0.05, ** p < 0.01. P-values were calculated using an unpaired, two sample t-test.

**Supplementary Figure 2:**
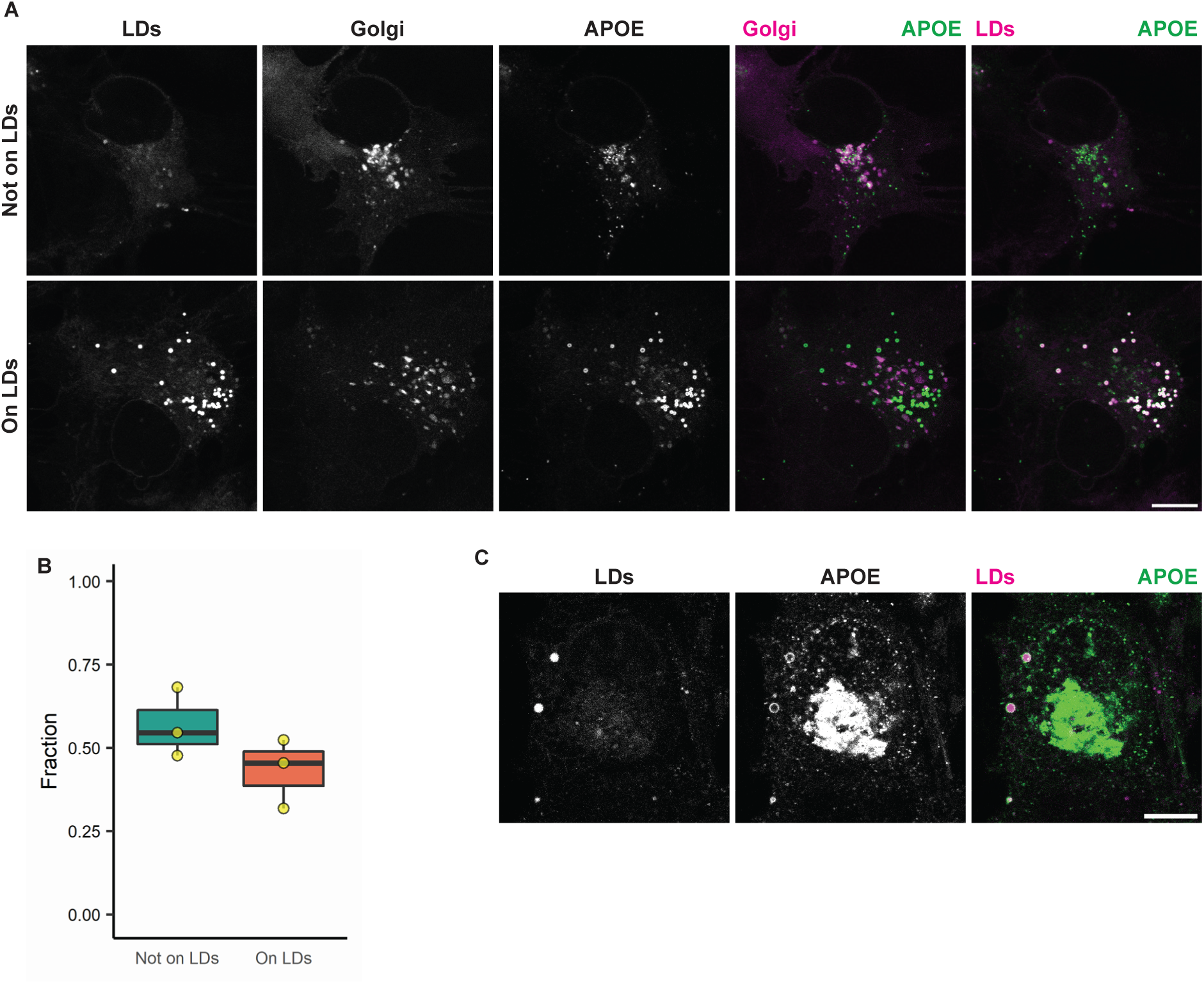
Exogenous APOE targets LDs in primary rat cortical astrocytes and U-2 OS cells. (**A**) Representative confocal slices of primary rat cortical astrocytes grown in astrocyte growth media (AGM) without OA supplementation, transfected with human APOE3-mEm and the Golgi marker mApple-SiT, and labelled for LDs with BODIPY 665/676. In the merged images, APOE is in green and the Golgi or LDs are in magenta as indicated. In primary astrocytes grown in AGM, APOE coats LDs in a subset of cells even in the absence of OA treatment. In cells where APOE does not coat LDs, APOE localizes to the Golgi. Scale bar, 10 µm. **(B)** Quantification of the fraction of transfected primary astrocytes with APOE on LDs vs. APOE not on LDs. Data was collected from 3 independent experiments. Each data point is the fraction of cells with APOE on LDs or APOE not on LDs from 44-63 randomly selected cells in one independent experiment. **(C)** Representative confocal slices of U-2 OS cells grown in CM without OA supplementation, transfected with APOE3-mEm, and labelled for LDs with BODIPY 667/676. Scale bar, 10 µm.

**Supplementary Figure 3:**
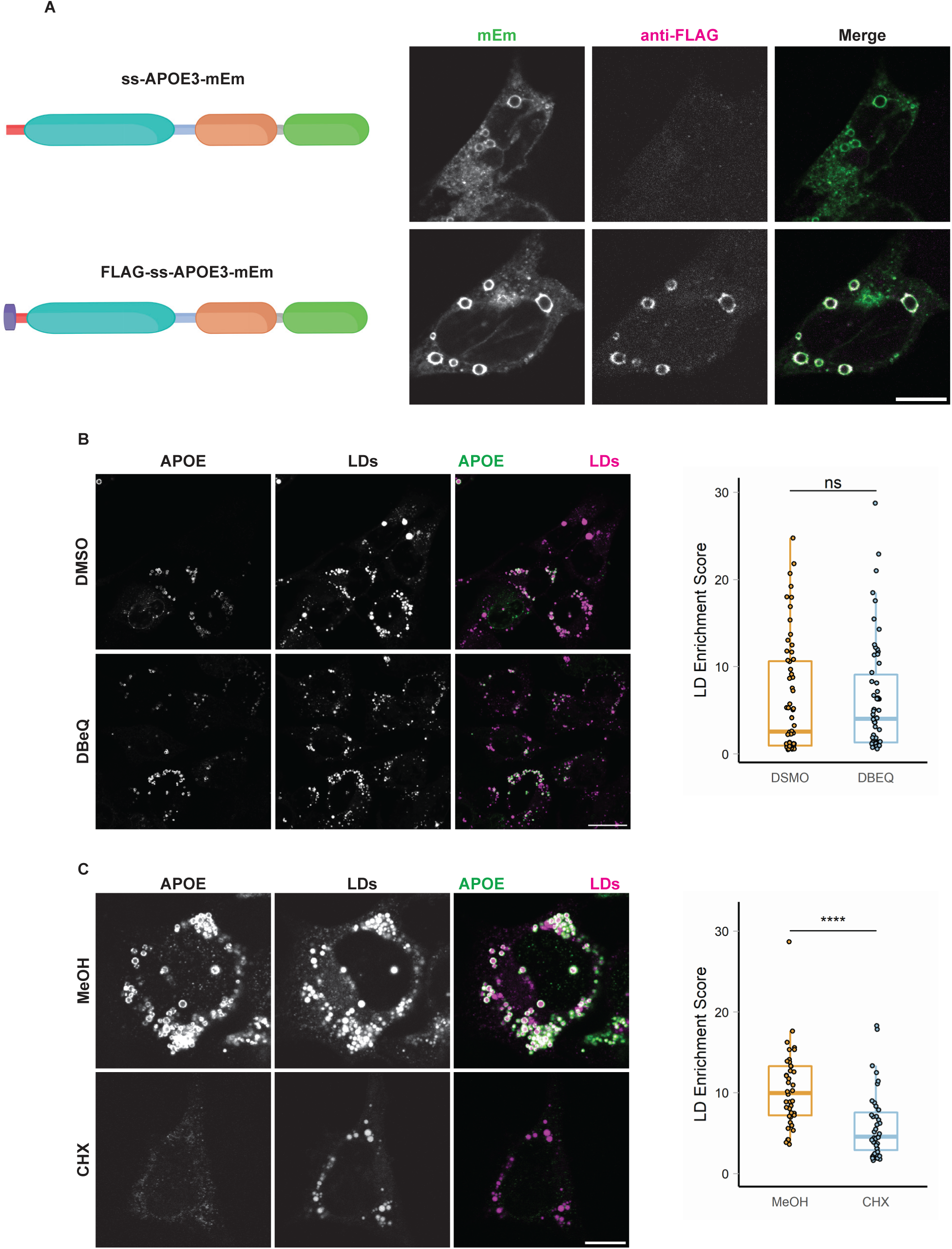
LD-associated APOE retains its signal peptide and is not retrotranslocated from the ER. (**A**) Representative confocal slices of TRAE3-H cells transfected with APOE3-mEm or FLAG-SS-APOE3-mEm and treated with 400 µM OA for 5 hours. Cells were then fixed and stained for the FLAG tag with an anti-FLAG antibody. Fluorescence signal on the surface of LDs is positive for both FLAG and mEmerald, indicating that LD-associated APOE retains its N-terminal signal peptide. By contrast, APOE in the secretory pathway is mEm positive but does not stain for FLAG, indicating that the pool of APOE in the secretory pathway is properly processed. **(B)** Representative confocal slices of TRAE3-H treated with 400 µM OA for 5 hours together with 0.1% DMSO vehicle or 10 µM DBeQ. Cells were then fixed and stained for endogenous APOE with an anti-APOE antibody and labelled for LDs with BODIPY 493/507. In the merged image, APOE is in green and LDs are in magenta. The LD enrichment fraction was calculated as described in Figure 1. There is no significant difference in LD enrichment upon DBeQ-treatment, indicating that p97-dependent retrotranslocation is not required for LD-targeting of APOE. N = 50 cells per condition. Data was collected and pooled from three biologically independent experiments. Scale bar, 10 µm. ns, p>0.05. **(C)** Representative confocal slices of TRAE3-H slices treated with 400 µM OA for 5 hours together with 0.1% MeOH vehicle or 100 µg/mL cycloheximide. Cells were then fixed and stained for endogenous APOE with an anti-APOE antibody and labelled for LDs with BODIPY 493/507. In the merged image, APOE is in green and LDs are in magenta. The LD enrichment fraction was calculated as described in Figure 1. There is a significant reduction in APOE on LDs upon cycloheximide treatment, suggesting that LD-associated APOE is newly translated. N = 50 cells per condition. Scale bar, 10 µm. ns, p>0.05. P-values were calculated using a Wilcox rank sum test.

**Supplementary Figure 4:**
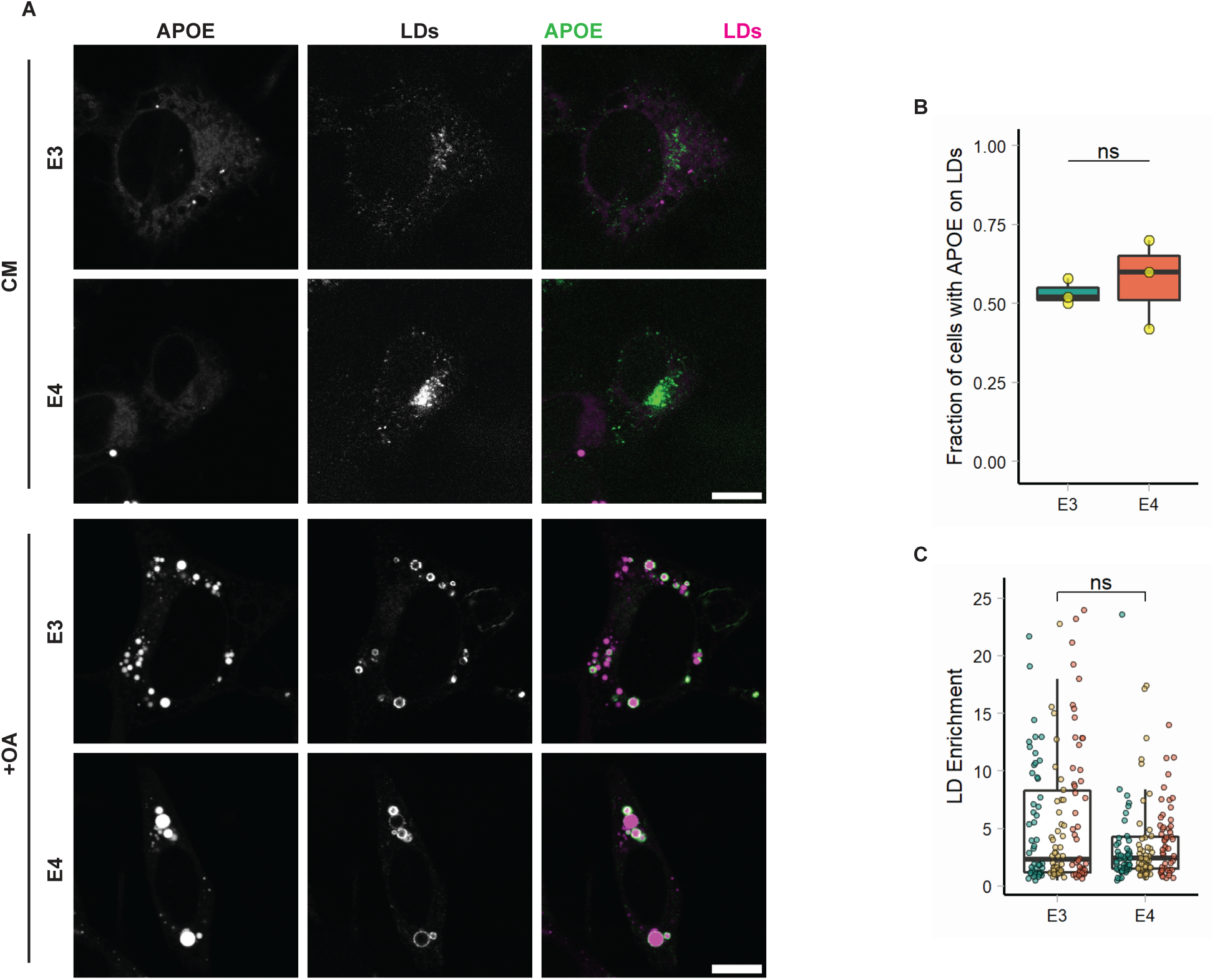
Targeting to LDs is unaffected in APOE4. (**A**) Representative confocal images of TRAE3-H cells or TRAE4-H cells with or without 5 hours of OA loading. Cells were fixed and stained for endogenous APOE with anti-APOE antibody and for LDs with BODIPY 493/507. In merged images, APOE is in green and LDs are in magenta. Scale bars, 10 µm. CM, complete media. +OA, 400 µM OA for 5 hours. **(B)** Fraction of TRAE3-H or TRAE4-H cells with APOE on the surface of LDs after 5 hours of OA. Each data point is the fraction of cells from 10 random fields of view with APOE on LDs in one experiment. APOE localization to LDs was determined qualitatively. N = 3 biologically independent experiments with 50 cells per experiment. ns, p > 0.05. P-value was calculated using an unpaired, two-sample t-test. **(C)** LD enrichment fraction of TRAE3-H or TRAE4-H cells treated with OA for 5 hours. LD enrichment fraction was calculated as described in Figure 1. N = 50 cells per condition and experiment. Each data point represents one cell, and each color represents data collected from a separate, independent experiment. These are the same cells used in B, but LD enrichment was measured using an unbiased quantitative method rather than being assessed qualitatively. ns, p>0.05. P-value was calculated using a clustered Wilcox rank sum test via the Rosner-Glynn-Lee method.

**Video 1: Partial and full rings of APOE surrounding LDs contacting the ER**

Airyscan live-cell imaging of TRAE3-H cells transfected with the ER lumen marker TagBFP2-KDEL (cyan) and APOE3-mEm (yellow), stained for LDs (magenta) with BODIPY 665/676, and treated with 400 µM OA for 4 hours. The video shows both full and half rings of APOE on the surface of LDs and at ER-LD contact sites. Scale bar: 2 µm. Corresponds to images shown in Fig. 3A. Cells were imaged every 7.9 seconds for 80 frames. Video plays at 10 frames per second.

**Video 2: Full rings of APOE surrounding LDs contacting the ER**

Airyscan live-cell imaging of TRAE3-H cells transfected with the ER lumen marker TagBFP2-KDEL (cyan) and APOE3-mEm (yellow), stained for LDs (magenta) with BODIPY 665/676, and treated with 400 µM OA for 4 hours. The video shows full rings of APOE on the surface of LDs and at ER-LD contact sites. Scale bar: 1 µm. Corresponds to images shown in Fig. 3A. Cells were imaged every 4.9 seconds for 29 frames. Video plays at 10 frames per second.

**Video 3: FRAP of LD-associated APOE during an OA pulse**

Confocal FRAP movies of primary astrocytes transfected with APOE3-mEm (green), stained for LDs with BODIPY 665/676 (magenta), and treated with 200 µM OA for 4 hours. LD-associated APOE rapidly recovers when bleached during an OA pulse. Scale bar: 1 µm. Corresponds to images shown in Fig. 3C. Cells were imaged every second for 96 frames. Video plays at 5 frames per second.

**Video 4: FRAP of LD-associated APOE after an OA pulse-chase**

Confocal FRAP movies of primary astrocytes transfected with APOE3-mEm (green), stained for LDs with BODIPY 665/676 (magenta), and treated with 200 µM OA for 4 hours followed by a 2-hour chase in complete media. LD-associated APOE recovers very slowly when bleached after an OA pulse-chase. Scale bar: 1 µm. Corresponds to images shown in Fig. 3C. Cells were imaged every second for 96 frames. Video plays at 5 frames per second.

